# Widespread introgression across a phylogeny of 155 *Drosophila* genomes

**DOI:** 10.1101/2020.12.14.422758

**Authors:** Anton Suvorov, Bernard Y. Kim, Jeremy Wang, Ellie E. Armstrong, David Peede, Emmanuel R. R. D’Agostino, Donald K. Price, Peter Wadell, Michael Lang, Virginie Courtier-Orgogozo, Jean R. David, Dmitri Petrov, Daniel R. Matute, Daniel R. Schrider, Aaron A. Comeault

## Abstract

Genome-scale sequence data have invigorated the study of hybridization and introgression, particularly in animals. However, outside of a few notable cases, we lack systematic tests for introgression at a larger phylogenetic scale across entire clades. Here we leverage 155 genome assemblies, from 149 species, to generate a fossil-calibrated phylogeny and conduct multilocus tests for introgression across nine monophyletic radiations within the genus *Drosophila*. Using complementary phylogenomic approaches, we identify widespread introgression across the evolutionary history of *Drosophila*. Mapping gene-tree discordance onto the phylogeny revealed that both ancient and recent introgression has occurred across most of the nine clades that we examined. Our results provide the first evidence of introgression occurring across the evolutionary history of *Drosophila* and highlight the need to continue to study the evolutionary consequences of hybridization and introgression in this genus and across the Tree of Life.

## INTRODUCTION

The extent of gene exchange in nature has remained one of the most hotly debated questions in speciation genetics. Genomic data have revealed that introgression is common across taxa, having been identified in major groups such as fungi^1–3^, vertebrates^4–7^, insects^8–10^, and angiosperms^11,12^. The evolutionary effects of introgression are diverse, and are determined by multiple ecological and genomic factors^13,14^. Once thought to be strictly deleterious, it has become increasingly clear that introgression can serve as a source of genetic variation used during local adaptation^15,16^ and adaptive radiation^17,18^. While our understanding of introgression as a widespread phenomenon has clearly improved, it remains unclear how often it occurs across taxa. Ideally, determining the frequency of introgression across the Tree of Life would leverage the signal from systematic analyses of clade-level genomic data without an *a priori* selection of taxa known to hybridize in nature.

At the phylogenetic scale, hybridization has typically been explored at relatively recent timescales. For example, studies of hybridization between cats (Felidae; 10-12 My; ~40 species^19^), butterflies (*Heliconius*; 10-15 My; 15 species^8^), cichlid fishes from the African rift lakes (0.5-10 My; ~27 species^18,20,21^), and wild tomatoes (*Solanum*; ~4 My; ~20 species^12^) all rejected a purely bifurcating phylogenetic history. In each of these systems introgression has occurred relatively recently, as the common ancestor for each species group occurred no more than 15 million years ago. However, there are also notable exceptions, and evidence for introgression has been found across much deeper phylogenetic timescales within vascular plants^11^ and primates^7^. In some species, there is also evidence that introgression has been a source of adaptive genetic variation that has helped drive adaptation (e.g. refs. 2,22–25). These results show how introgression has both (1) occurred in disparate taxonomic groups and (2) promoted adaptation and diversification in some. Notwithstanding key examples^4–7,11,12^, we still require systematic tests of introgression that use clade-level genomic data that spans both deep and shallow phylogenetic time to better understand introgression’s generality throughout evolution.

Species from the genus *Drosophila* remain one of the most powerful genetic systems to study animal evolution. Comparative analyses suggest that introgression might be common during speciation in the genus^26^. Genome scans of closely related drosophilid species have provided evidence of gene flow and introgression^9,10,27–32^. There is also evidence of contemporary hybridization^33–35^ and stable hybrid zones between a handful of species^36–38^. These examples of hybridization and introgression show that species boundaries can be porous but cannot be taken as *prima facie* evidence of the commonality of introgression. We still lack a systematic understanding of the relative frequency of hybridization and subsequent introgression across *Drosophila*. Here we analyze patterns of introgression across a phylogeny generated using 155 whole genomes derived from 149 species of *Drosophila*, and the genomes of four outgroup species. These *Drosophila* species span over 50 million years of evolution and include multiple samples from nine major radiations within the genus *Drosophila*. We used two different phylogenetic approaches to test whether introgression has occurred in each of these nine radiations. We found numerous instances of introgression across the evolutionary history of drosophilid flies, some mapping to early divergences within clades up to 20-25 Mya. Our results provide a taxonomically unbiased estimate of the prevalence of introgression at a macroevolutionary scale. Despite few known observations of current hybridization in nature, introgression appears to be a widespread phenomenon across the phylogeny of *Drosophila*.

## RESULTS

### A high-confidence phylogeny of 155 *Drosophila* genomes

We first used genome-scale sequence data to infer phylogenetic relationships among species in our data set. To achieve this, we annotated and generated multiple sequence alignments for 2,791 Benchmarking Universal Single-Copy Orthologs (BUSCOs; v3^39,40^) across 155 independently assembled *Drosophila* genomes together with four outgroups (3 additional species from Drosophilidae and *Anopheles gambiae*). We used these alignments, totalling 8,187,056 nucleotide positions, and fossil calibrations to reconstruct a fossil-calibrated tree of *Drosophila* evolutionary history. Note that the inclusion of *Anopheles* as an outgroup allowed us to include a fossil of *Grauvogelia*, the oldest known dipteran, in our fossil calibration analysis, along with several *Drosophilidae* fossils and/or geological information (i.e., formation of the Hawaiian Islands; Data S1).

Our phylogenetic analyses (see Method Details for details) using both maximum-likelihood (ML using the `IQ-TREE` package) and gene tree coalescent-based (`ASTRAL`) approaches with DNA data revealed well-supported relationships among nearly all species within our dataset. Phylogenies inferred using these two approaches only differed in three relationships (Figure S1): (i) *D. villosipedis* was either recovered as sister species to *D. limitata* + *D. ochracea* (ML topology) or as a sister to *D. limitata* + *D. ochracea* + *D. murphyi* + *D. sproati* (ASTRAL topology); (ii) *D. vulcana* and *D. seguyi* form monophyletic lineage sister to the *D. nikananu* + *D. spp. aff. chauvacae* + *D. burlai* + *D. bocqueti* + *D. bakoue* clade (ML topology) or have paraphyletic relationships where *D. vulcana* is sister to the *D. nikananu* + *D. spp. aff. chauvacae* + *D. burlai* + *D. bocqueti* + *D. bakoue* clade (ASTRAL topology) ; (iii) *D. simulans* was recovered as sister either to *D. mauritiana* (ML topology) or *D. sechellia* (ASTRAL topology, the latter of which is perhaps more likely to be the true species tree according to an analysis examining low-recombining regions, which are less prone to ILS^41^. The nodal supports were consistently high across both ML (Ultrafast bootstrap (UFBoot) = 100, an approximate likelihood ratio test with the nonparametric Shimodaira–Hasegawa correction (SH-aLRT) = 100, a Bayesian-like transformation of aLRT (aBayes) = 1) and ASTRAL (Local posterior probability (LPP) = 1) topologies with the exception of *D. limitata* + *D. ochracea* + *D. villosipedis* (UFBoot = 9, SH-aLRT = 81, aBayes = 1) and *D. carrolli* + *D. rhopaloa* + *D. kurseongensis* (UFBoot = 81.2, SH-aLRT = 81, aBayes = 1) on the ML tree, and *D. limitata* + *D. ochracea* + *D. murphyi* + *D. sproati* (LPP = 0.97) and *D. sulfugaster bilimbata*+ *D. sulfugaster sulfurigaster* (LPP = 0.69) on the ASTRAL tree. Thus, the phylogeny we report here is the first of the genus *Drosophila* with almost all nodes resolved with high confidence—recent estimates of the *Drosophila* phylogeny lacked strong support throughout all tree depth levels^42–44^.

Erroneous orthology inference as well as misalignment can impede accurate phylogenetic inference and create artificially long branches^45^. Repeating our ASTRAL analysis after removing outlier long branches via `TreeShrink`^45^ resulted in an identical tree topology with the aforementioned ASTRAL tree (Figure S1). Furthermore, an ML topology estimated from the dataset with more closely related outgroup species (see Method Details) results in an identical topology with the aforementioned ML tree (Figure S1). The inferred phylogeny from the protein supermatrix showed only four incongruencies with the phylogeny that was inferred from DNA data (Figure S1): (i) *D. villosipedis* was recovered as a sister species to *D. limitata* + *D. ochracea* + *D. murphyi* + *D. sproati;* (ii) *D. watanabei* + *D. punjabiensis* is sister to the clade containing *D. bakoue* and *D. jambulina;* (iii) *D. vulcana* and *D. seguyi* show paraphyletic relationships; (iv) *Z. vittiger* and *Z. lachaisei* show sister species relationships. We performed further assessment of nodal support with Quartet Sampling^11^, using the Quartet Concordance (QC) and Quartet Differential (QD) scores to identify quartet-tree species-tree discordance (Method Details). At some nodes, an appreciable fraction of quartets disagreed with our inferred species tree topology (QC < 1), and in most of these cases this discordance was skewed toward one of the two possible alternative topologies (i.e. QD < 1 but > 0) as is consistent with introgression. We formally explore this pattern below.

In order to estimate divergence times across the *Drosophila* phylogeny, we developed five calibration schemes (A, B, C, D and “Russo”; Data S1) used in `MCMCtree`^46^ and one scheme based on the Fossilized Birth-Death (FBD) process^47^ used in `BEAST2`^48^ (BEAST2 FBD; Data S1). Overall, four of the five `MCMCtree` schemes yielded nearly identical age estimates with narrow 95 % credible intervals (CI), whereas scheme “Russo” (a fossil calibration strategy closely matching that from^43^) showed slightly older estimates (Figure S2) with notably wider 95% CIs. Throughout this manuscript we use the time estimates obtained with scheme A. This calibration analysis estimated that extant members of the genus *Drosophila* branched off from the other Drosophilidae (*Leucophenga, Scaptodrosophila* and *Chymomyza*) ~53 Mya (95% CI: 50 - 56.6 Mya) during the Eocene Epoch of the Paleogene Period (Figure 1). The same analysis inferred that the split between the two major lineages within *Drosophila—*the subgenera of *Sophophora* and *Drosophila*—occurred ~47 Mya (95% CI: 43.9-49.9 Mya; Figure 1; “A” and “B” clades, respectively); previously published estimates of this time include ~32 Mya (95% CI: 25–40 Mya^49^), ~63 Mya (95% CI: 39–87 Mya^50^), and ~56 Mya (95% CI not available^43^). We also note that our divergence time estimates of the *Drosophila* subgenus (~34 Mya, 95% CI: 31.6 - 36.8 Mya; clades 6 through 9) are somewhat younger than ~40 Mya, a previous estimate reported in^51^, although the latter had fairly wide confidence intervals (95% CI: 33.4 - 47.6 Mya). On the other hand, divergence time estimates produced by the FBD scheme in `BEAST2` tend to be older especially for deeper nodes (Figure S2). Also, CIs estimated by `BEAST2` were wider than those from `MCMCTREE`. This can be explained by the fewer assumptions about fossil calibration placement and age prior specification for methods that rely on the FBD process. Additionally, we note that not all parameters of the BEAST2 FBD calibration scheme converged (i.e., effective sample size < 100) even after 6 × 10^8^ MCMC generations. Thus, the lack of a thorough fossil record within *Drosophila* makes it difficult to accurately and precisely estimate divergence times, and point estimates of divergence times should be interpreted with caution.

**Fiure 1.**
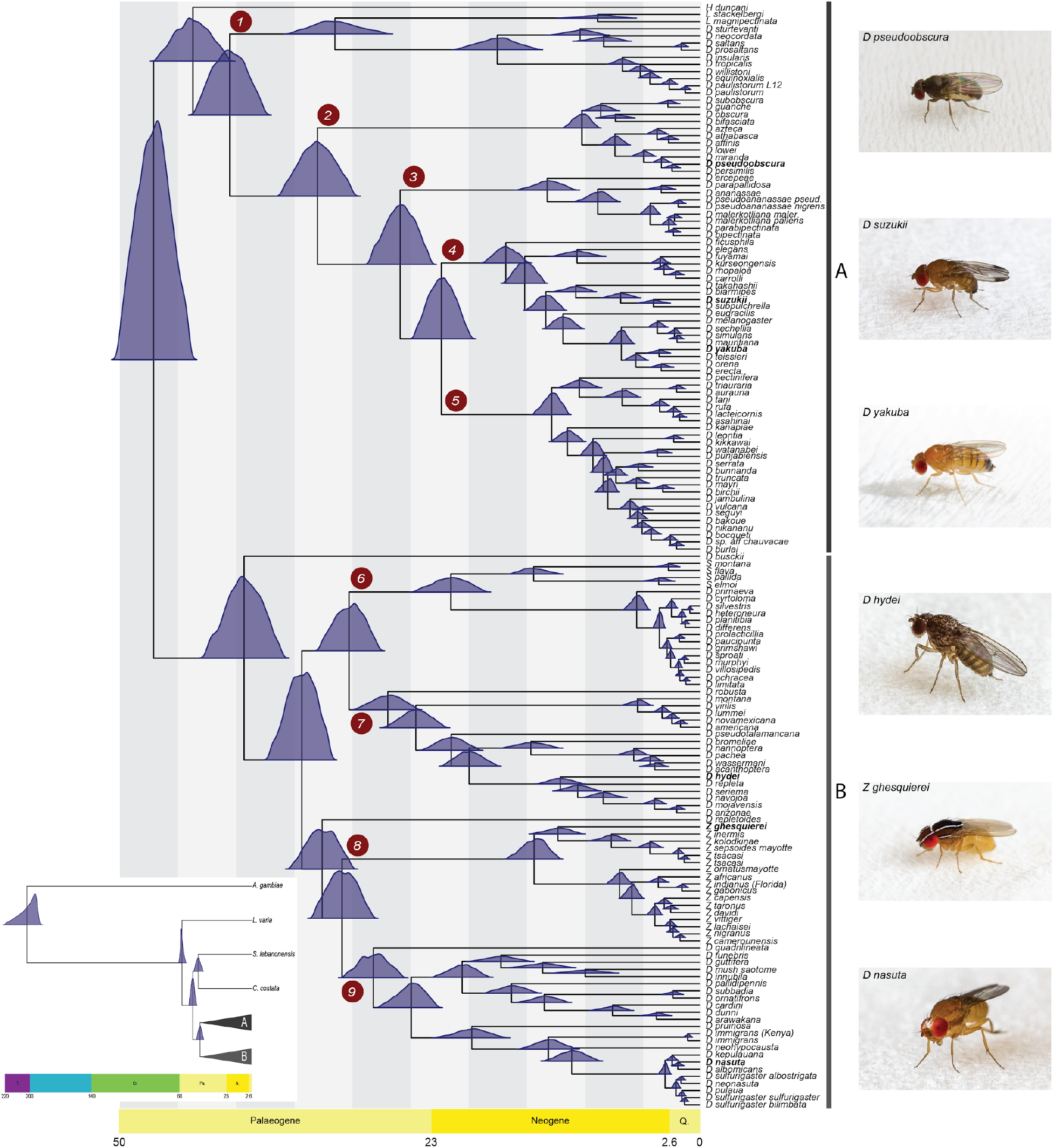
Fossil calibrated maximum likelihood phylogenetic tree of the genus *Drosophila* inferred from a supermatrix of 2,791 BUSCO loci (total of 8,187,056 sites). The blue distributions at each divergence point on the tree represent nodal age posterior probabilities from MCMCTree. *Grauvogelia* and *Oligophryne* fossils were used to set priors on the age of the root of the tree, *Phytomyzites* and *Electrophortica succini* were used for priors for the root of the Drosophilidae family, and *Electrophortica succini* and *Scaptomyza dominicana* were used to set priors for the crown group “*Scaptomyza”*, i.e. Most Recent Common Ancestor (MRCA) node of the *Scaptomyza* species (scheme A; Data S1). The numbered red circles denote clades for which analyses of introgression were performed. Inset: the phylogenetic and temporal relationships between our distant outgroup *Anopheles gambiae*, more closely related outgroup species of Drosophilidae (*Leucophenga varia*, *Scaptodrosophila lebanonensis* and *Chymomyza costata*), and the *Drosophila* genus. A and B denote the two inferred major groups within *Drosophila*.

### Widespread signatures of introgression across the *Drosophila* phylogeny

To assess the prevalence of introgression across the *Drosophila* tree, we subdivided species into nine monophyletic lineages (herein referred to as clades 1 through 9; Figure 1) and tested for introgression within each clade. These clades correspond to the deepest divergences within the genus, with most having an MRCA during the Paleogene. Clades 4 and 5 are the two exceptions, splitting from an MRCA later in the Neogene. Within each of the nine clades, the MRCA of all sampled genomes ranged from ~10 Mya (Figure 1; clade 2) to ~32 Mya (Figure 1; clade 1). We note that *Hirtodrosophila duncani, Drosophila busckii* and *Drosophila repletoides* were not included in these clade assignments as each of these species was the only sampled descendent of a deep lineage; additional taxon sampling is required to assign them to specific monophyletic species groups that could be tested for introgression.

We tested for introgression within each of these nine clades using two complementary phylogenomic methods that rely on the counts of gene trees inferred from the BUSCO loci that are discordant with the inferred trees (hereafter referred to as the discordant-count test or DCT) and the distribution of branch lengths for discordant gene trees (hereafter termed the branch-length test or BLT), respectively, among rooted triplets of taxa (Figure 2). These methods leverage information contained across a set of gene trees to differentiate patterns of discordance that are consistent with introgression from those that can be explained by incomplete lineage sorting alone (see Method Details). We found at least one pair of species with evidence of introgression in 7 of the 9 clades according to both DCT and BLT (i.e. the same pair of species showed evidence for introgression that was significant for both tests in the same triplet at an FDR-corrected *P*-value threshold of 0.05). In clades 1 and 3 there were no species pairs for which the DCT and BLT were significant in the same triplet and both suggest the same introgressing species pair (Data S2). However, both clades had several pairs that were significant according to one test or the other (Data S2). We found even stronger support for introgression using two existing software methods: QuIBL (Data S2), which examines the branch-length distributions of all three gene tree topologies for a triplet^8^, and HyDe (Data S2), which tests for introgression by counting quartet site patterns^52^. Specifically, QuIBL detected introgression in 120 of 152 (78.9%) of species pairs detected by both DCT and BLT, as well as 894 additional species pairs not detected by DCT-BLT; we note that BLT and QuIBL approaches are not fully independent, since they both utilize branch-length information. Similarly, HyDe detected introgression in 142 of 152 (93.4%) of species pairs detected by both DCT and BLT, and 898 additional species pairs (the results of HyDe were not qualitatively affected if a more distantly related outgroup, i.e. *Anopheles gambiae*, was selected, see Data S2). However, we focus here on the intersection between DCT and BLT methods (after correcting each for multiple testing), as this provides a more conservative estimate of the extent of introgression. Supporting this claim, we applied these tests to a gene tree dataset simulated under high levels of ILS^53^ and observed low false positive rates: 0.054 for DCT, 0.089 for BLT, and 0.009 for their intersection.

**Figure 2.**
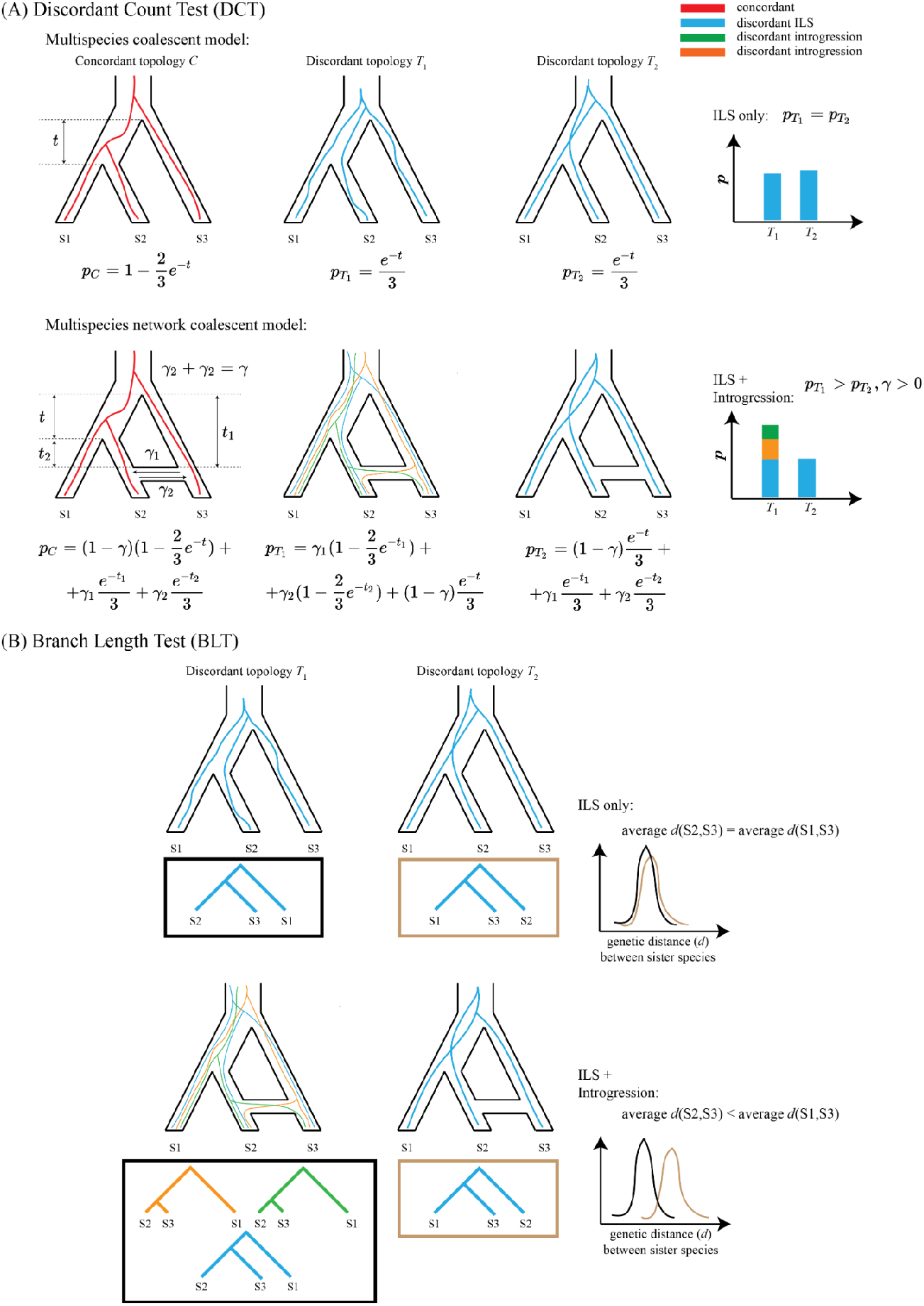
Overview of the phylogenomic approaches used to detect introgression. (A) The Discordant Count Test (DCT) identifies introgression where a given triplet within the tree shows an excess of gene trees that support one of the two possible divergent topologies. Note that concordant gene trees and corresponding probabilities are included for completeness, although these are not used by our test. (B) The Branch Length Test (BLT) identifies introgression where branch lengths of gene trees that support introgression are shorter than branch lengths of those that support the species tree and the less frequent divergent topology (i.e., the discordant topology putatively due to ILS).

We carried out several analyses to assess the robustness of our results to data quality and evolutionary rate. First, to assess the effect of alignment length we performed BLT and DCT analyses on gene trees that excluded alignments with fewer than 1,000 sites. We found that ~ 84%, ~94% and ~80% of introgressing species pairs that were identified by BLT, DCT, and their intersection, respectively, remained significant after filtering out these short alignments (Data S2). Second, we explored whether the rapid karyotype evolution^54^ observed in the *obscura* group (our clade 2) may impact introgression inference. To that end, we excluded loci that did not belong to the same Muller element within each analyzed triplet by BLT and DCT. This filtering scheme had a minor impact on introgression estimation with ~70%, ~89% and ~76% introgressing taxon pairs identified by BLT, DCT and their intersection being identified after filtering out loci that are found on different Muller elements in species within this clade (Data S2). More importantly, this filtering had no impact on the introgression events discussed below and shown in Figure 2—the exact same events were inferred for clade 2. Third, we investigated the effects of evolutionary rate (as measured by *d*_N_/*d*_S_) heterogeneity across branches on introgression inference. For each triplet tested by BLT and DCT we excluded gene trees with *d*_N_/*d*_S_ > 0.53, which corresponds to the 5% critical value of *d*_N_/*d*_S_ distribution across all the clades and gene trees. Overall, ~69%, ~82% and ~54% of our original number of introgressing taxon pairs were identified after *d*_N_/*d*_S_ filtering by the BLT, DCT and their intersection, respectively (Data S2). Importantly, a large number of genes are removed when applying this filter because only one branch within the portion of the gene tree relevant to the triplet must exceed the critical value of *d*_N_/*d*_S_ to result in the entire gene tree’s removal. We therefore asked to what extent this reduced fraction of introgressing taxon pairs is a consequence of reduced power due to the reduction in the number of gene trees. We found that randomly subsampling gene trees without respect to *d*_N_/*d*_S_ value can affect introgression inferences in a similar fashion: on average ~76%, ~85% and ~65% introgressing taxon pairs were identified by BLT, DCT and their intersection, respectively, after randomly removing the same number of genes removed by our *d*_N_/*d*_S_ filter (see Method Details). Thus, although we don’t rule out the possibility that evolutionary rate heterogeneity may influence our DCT-BLT analysis, or that the persistence of introgressed alleles may be correlated with a gene’s evolutionary rate, this result shows that our estimates of gene flow are not being driven primarily by genes evolving under the least amount of selective constraint and/or the greatest amount of positive selection. We also repeated our BLT and DCT analyses using a gene tree set with potentially misaligned sequences removed via `TreeShrink` and obtained results largely concordant with other methods as shown in Data S2. However, we notice several exceptions: in clades 5 and 7 the number of species pairs with at least one triplet that is significant according to both the DCT and BLT methods is markedly higher after running `TreeShrink`, largely due to an increase in significant DCT results. In addition, for Hawaiian drosophilids (clade 6) we find no introgression based on the overlap between BLT and DCT.

The number of species pairs that show evidence of introgression in our initial DCT-BLT analysis is not equivalent to the number of independent introgression events among *Drosophila* species. This is because gene flow in the distant past can leave evidence of introgression in multiple contemporary species pairs. For example, we found evidence for introgression between *D. robusta* and all five species within the *D. americana-D. montana* group (see clade 7 in Figure 3). Rather than five independent instances of introgression between species, this pattern could reflect introgression between ancestral taxa that subsequently diverged into the contemporary species. More generally, cases where multiple introgressing species pairs each shared the same MRCA may be more parsimoniously explained by a single ancestral introgression event between the branches that coalesce at this node, while those involving only a single species pair may have resulted from introgression between the extant species pair (Data S2). Another example of the former can be seen in clade 6 where the evidence suggests introgression occurred between the Hawaiian *Scaptomyza* and *Drosophila* (Figure S3) that are estimated to have diverged from each other more than 20 Mya. This ancient introgression may have occurred prior to the formation of Kauai island ~5 Mya which is now the oldest high island with extant species in these two groups^55,56^.

**Figure 3.**
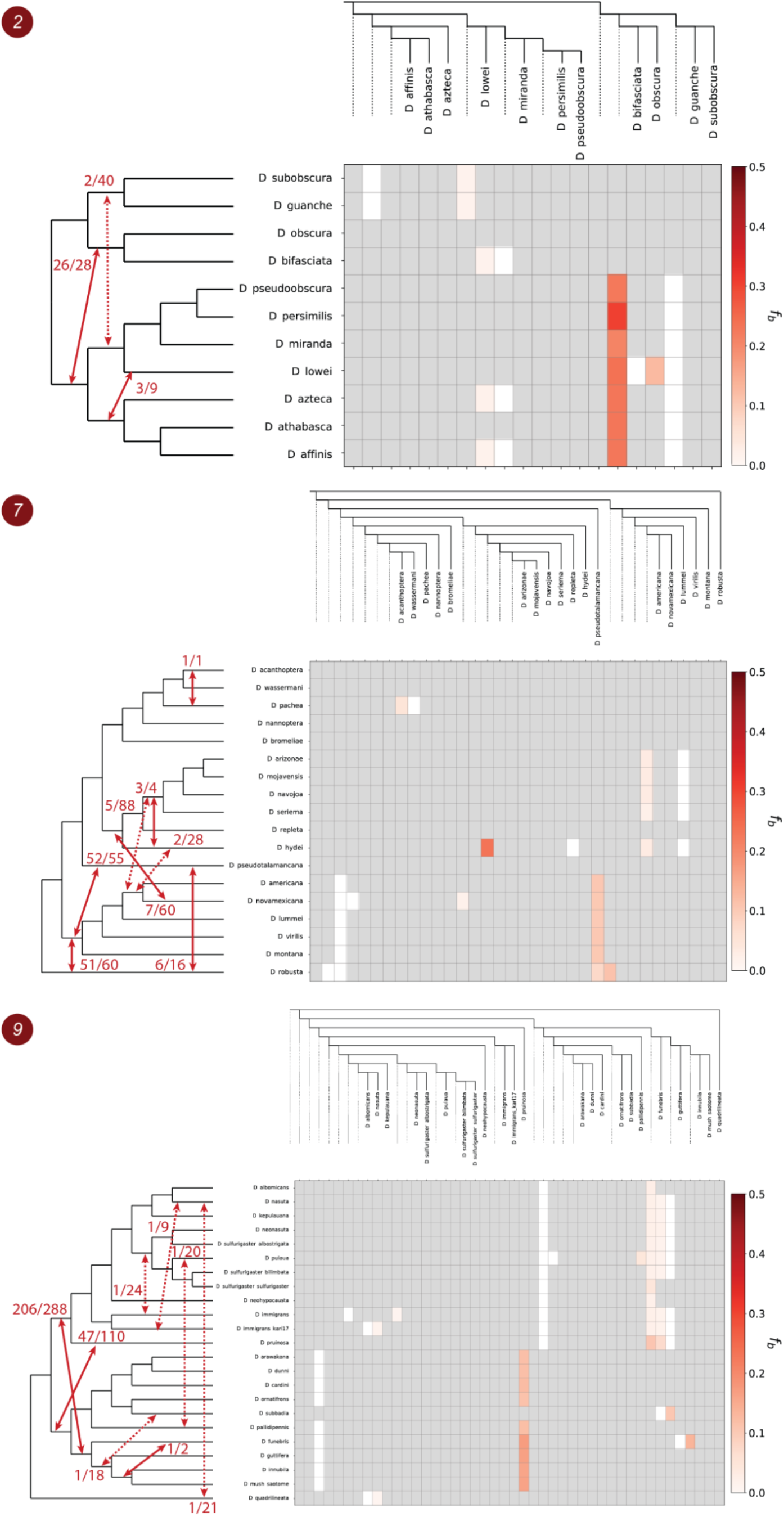
Patterns of introgression inferred for the monophyletic clades 2, 7 and 9. The matrix shows inferred introgression proportions as estimated from gene tree counts for the introgressed species pairs (Method Details), and then mapped to internal branches using the *f*-branch method^20^. The expanded tree at the top of each matrix shows both the terminal as well as ancestral branches. The tree on the left side of each matrix represents species relationships with mapped introgression events (red arrows) derived from the corresponding *f*-branch matrix (Method Details). The fractions next to each arrow represent the number of triplets that support a specific introgression event by both DCT and BLT divided by the total number of triplets that could have detected the introgression event. Dashed arrows represent introgression events with low support (triplet support ratio < 10%).

To summarize our DCT-BLT results and estimate both the number of introgression events and the proportion of the genome that introgressed during those events (*γ*) we adapted the *f*-branch heuristic^20^ (implemented in `Dsuite`^57^; Method Details). Summed across all clades, our *f*-branch results suggest that at least 30 introgression events are required to explain our DCT-BLT results (Figure 3 and Figure S2). Clades 2, 4, 6, 7 and 9 showed the strongest evidence of introgression, in terms of both the total number of DCT-BLT significant triplets and *γ* estimates from `Dsuite` that support those events (Table 1). For example, in clade 2 Dsuite suggests an ancestral introgression event between the branch leading to *D. obscura* and *D. bifasciata* and the branch that leads to the clade containing *D. pseudoobscura* and *D. affinis*. Furthermore, this particular signal is characterized by a large fraction of introgressed genetic material (*γ* = 0.237, Table 1) and by the large number of triplets that are significant according to both DCT and BLT (26 out of 28 total triplets that could detect this event are significant according to both tests). We stress that both our methods used to detect introgression (DCT and BLT) and our approaches for counting introgression events (*f*-branch) are conservative, and thus the true number of events could be substantially greater, as suggested by our analyses using QuIBL and HyDe. Regardless of the method used, careful examination of results in Data S2, Figure 3 and Figure S2 reveals that deep introgression events are clearly the best explanation for some of our patterns (e.g. the case from clade 7 involving *D. robusta* described above), although more recent events may have occurred as well (e.g. between *D. pachea* and *D. acanthoptera*; Data S2, clade 7).

**Table 1.**
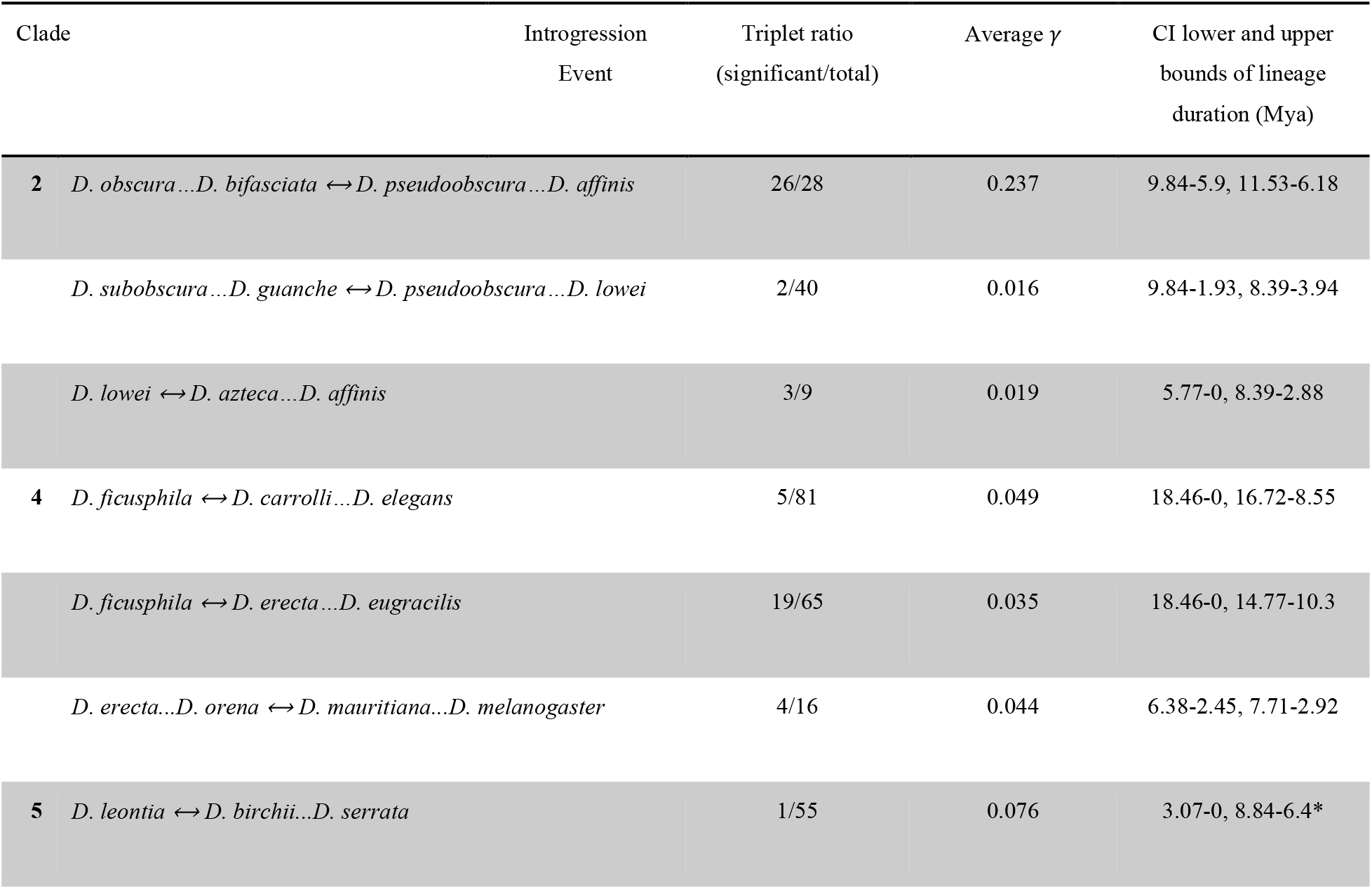
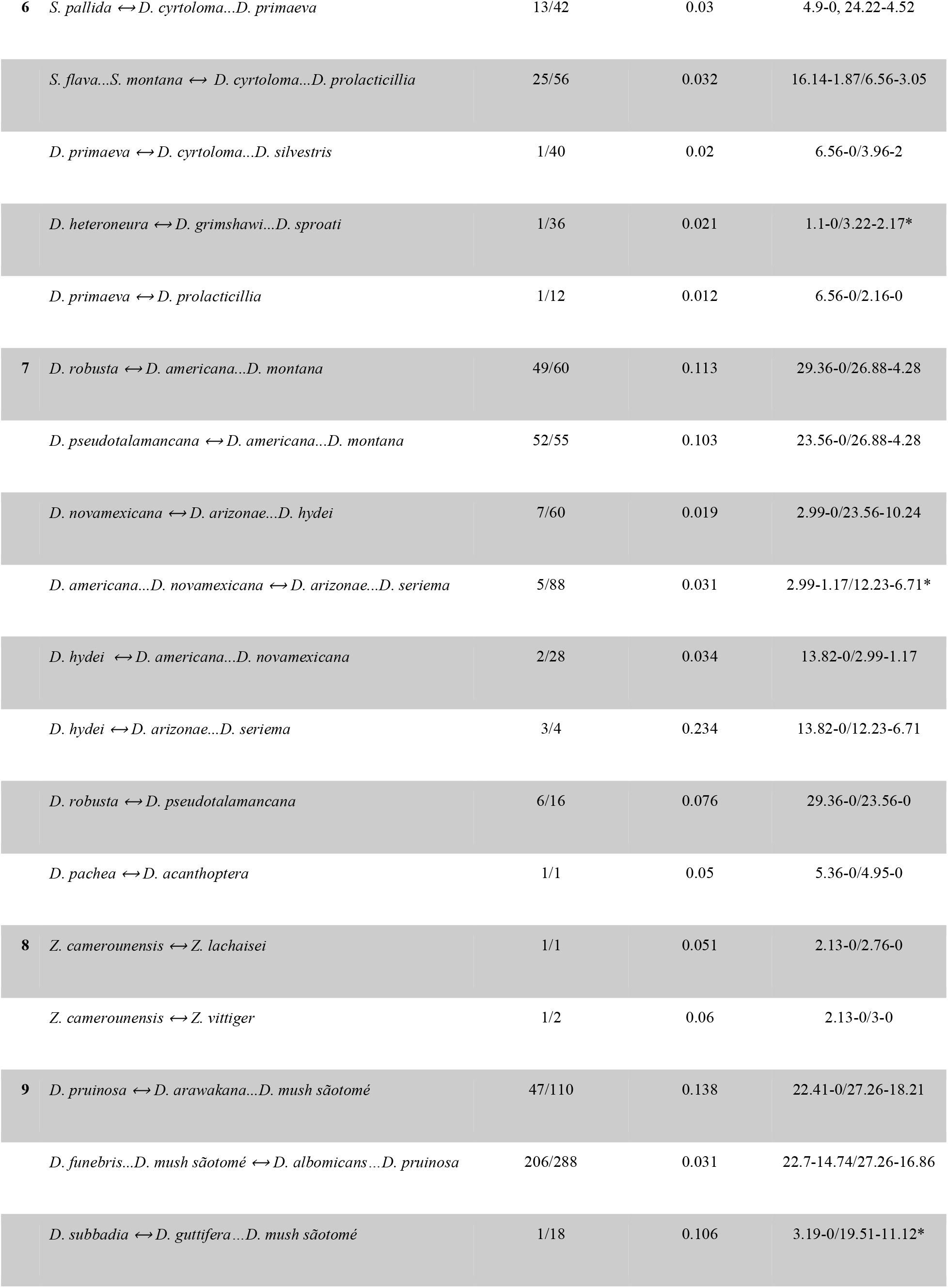
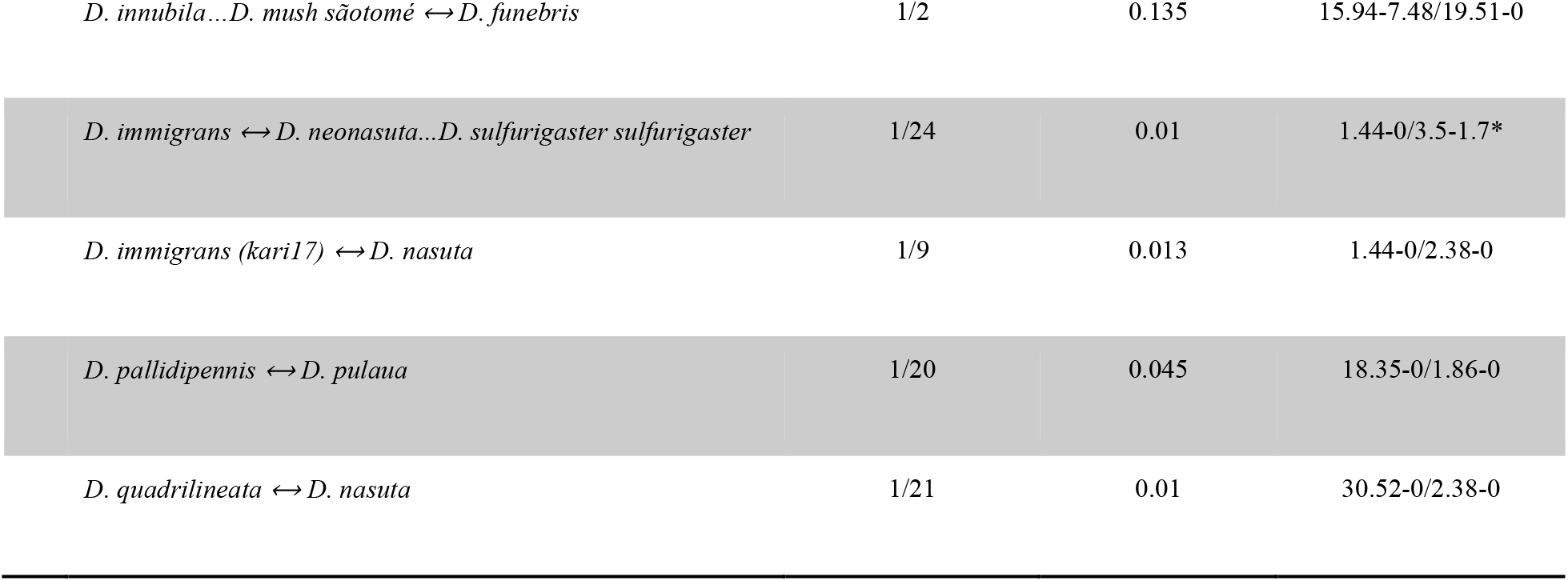
Placements, support and timing of introgression events across the *Drosophila* phylogeny. Putative introgression events (↔) are specified between different clades indicated by the pair of species in that clade with the oldest MRCA. The triplet ratio shows the number of significant and non-significant triplets according to DCT-BLT. The average *γ* was obtained from the *f*-branch results. The durations of the two introgressing lineages are represented by predicted lower and upper boundaries of credible intervals (95% CIs) estimated by MCMCTree using calibration scheme A. * indicates introgressing lineages with no time overlap (according to 95% CIs).

| Clade | Introgression Event | Triplet ratio (significant/total) | Average $\gamma$ | CI lower and upper bounds of lineage duration (Mya) |
| --- | --- | --- | --- | --- |
| 2 | *D. obscura*...*D. bifasciata* ↔ *D. pseudoobscura*...*D. affinis* | 26/28 | 0.237 | 9.84-5.9, 11.53-6.18 |
| | *D. subobscura*...*D. guanche* ↔ *D. pseudoobscura*...*D. lowei* | 2/40 | 0.016 | 9.84-1.93, 8.39-3.94 |
| | *D. lowei* ↔ *D. azteca*...*D. affinis* | 3/9 | 0.019 | 5.77-0, 8.39-2.88 |
| 4 | *D. ficusphila* ↔ *D. carrolli*...*D. elegans* | 5/81 | 0.049 | 18.46-0, 16.72-8.55 |
| | *D. ficusphila* ↔ *D. erecta*...*D. eugracilis* | 19/65 | 0.035 | 18.46-0, 14.77-10.3 |
| | *D. erecta*...*D. orena* ↔ *D. mauritiana*...*D. melanogaster* | 4/16 | 0.044 | 6.38-2.45, 7.71-2.92 |
| 5 | *D. leontia* ↔ *D. birchii*...*D. serrata* | 1/55 | 0.076 | 3.07-0, 8.84-6.4* |
| 6 | *S. pallida* ↔ *D. cyrtoloma*...*D. primaeva* | 13/42 | 0.03 | 4.9-0, 24.22-4.52 |
|  | *S. flava*...*S. montana* ↔ *D. cyrtoloma*...*D. prolacticillia* | 25/56 | 0.032 | 16.14-1.87/6.56-3.05 |
|  | *D. primaeva* ↔ *D. cyrtoloma*...*D. silvestris* | 1/40 | 0.02 | 6.56-0/3.96-2 |
|  | *D. heteroneura* ↔ *D. grimshawi*...*D. sproati* | 1/36 | 0.021 | 1.1-0/3.22-2.17* |
|  | *D. primaeva* ↔ *D. prolacticillia* | 1/12 | 0.012 | 6.56-0/2.16-0 |
| 7 | *D. robusta* ↔ *D. americana*...*D. montana* | 49/60 | 0.113 | 29.36-0/26.88-4.28 |
|  | *D. pseudotalamancana* ↔ *D. americana*...*D. montana* | 52/55 | 0.103 | 23.56-0/26.88-4.28 |
|  | *D. novamexicana* ↔ *D. arizonae*...*D. hydei* | 7/60 | 0.019 | 2.99-0/23.56-10.24 |
|  | *D. americana*...*D. novamexicana* ↔ *D. arizonae*...*D. seriema* | 5/88 | 0.031 | 2.99-1.17/12.23-6.71* |
|  | *D. hydei* ↔ *D. americana*...*D. novamexicana* | 2/28 | 0.034 | 13.82-0/2.99-1.17 |
|  | *D. hydei* ↔ *D. arizonae*...*D. seriema* | 3/4 | 0.234 | 13.82-0/12.23-6.71 |
|  | *D. robusta* ↔ *D. pseudotalamancana* | 6/16 | 0.076 | 29.36-0/23.56-0 |
|  | *D. pachea* ↔ *D. acanthoptera* | 1/1 | 0.05 | 5.36-0/4.95-0 |
| 8 | *Z. camerounensis* ↔ *Z. lachaisei* | 1/1 | 0.051 | 2.13-0/2.76-0 |
|  | *Z. camerounensis* ↔ *Z. vittiger* | 1/2 | 0.06 | 2.13-0/3-0 |
| 9 | *D. pruinosa* ↔ *D. arawakana*...*D. mush sãotomé* | 47/110 | 0.138 | 22.41-0/27.26-18.21 |
|  | *D. funebris*...*D. mush sãotomé* ↔ *D. albomicans*...*D. pruinosa* | 206/288 | 0.031 | 22.7-14.74/27.26-16.86 |
|  | *D. subbadia* ↔ *D. guttifera*...*D. mush sãotomé* | 1/18 | 0.106 | 3.19-0/19.51-11.12* |

**Table 1.** Placements, support and timing of introgression events across the *Drosophila* phylogeny.
|  |  |  |  |
| --- | --- | --- | --- |
| *D. innubila*...*D. mush* sãotomé ↔ *D. funebris* | 1/2 | 0.135 | 15.94-7.48/19.51-0 |
| *D. immigrans* ↔ *D. neonasuta*...*D. sulfurigaster sulfurigaster* | 1/24 | 0.01 | 1.44-0/3.5-1.7* |
| *D. immigrans (kari17)* ↔ *D. nasuta* | 1/9 | 0.013 | 1.44-0/2.38-0 |
| *D. pallidipennis* ↔ *D. pulaua* | 1/20 | 0.045 | 18.35-0/1.86-0 |
| *D. quadrilineata* ↔ *D. nasuta* | 1/21 | 0.01 | 30.52-0/2.38-0 |

We note that some scenarios of ancestral population structure could potentially result in differences in the number and branch lengths of gene trees with either discordant topology (discussed in Method Details). We therefore applied a more stringent version of the DCT-BLT that compares the branch lengths of the discordant topology with those of the concordant topology; this test will not be sensitive to ancestral population structure but could potentially produce many false negatives (Method Details). When applying this more stringent branch-length test to our data set, we find that of the 511 triplets significant according to our combined DCT-BLT test, 144 (28.1%) remain significant when imposing this much more stringent version of the BLT (again, after FDR correction). We then asked how many of the 30 introgression events shown in Figs. 3 and S3 were significant by this more stringent test for at least one triplet, finding that 13 of the 30 events (43.3%) are significant, including 11/17 (64.7%) of the most strongly supported events (those significant in at least 10% of triplets in our original analysis and shown in solid lines in Figs. 3 and S3). This result shows that the majority (~2/3) of our strongly supported putative introgression events are inconsistent with the phenomenon of ancestral population structure-produced false positives. Given that this test is highly conservative, we interpret this result as evidence that the vast majority of our detected introgression events are true positives rather than artifacts of population structure.

To complement our *f-*branch analysis, we also used PhyloNet^58,59^ to identify branches with the strongest signature of introgression in each of the nine monophyletic clades in our tree. Within each clade, we examined all possible network topologies produced by adding a single reticulation event to the species tree and determined which of the resulting phylogenetic networks produced the best likelihood score. We note that networks with more reticulation events would most likely exhibit a better fit to observed patterns of introgression but the biological interpretation of complex networks with multiple reticulations is more challenging; thus, we limited the analysis to a single reticulation event even though this will produce false negatives in clades with multiple gene flow events.

For all clades except clade 8, the networks with the highest likelihood scores from PhyloNet qualitatively agree with the inferred introgression patterns by the DCT-BLT results summarized by `Dsuite`: the best-supported position of a reticulation event inferred by PhyloNet tended to occur in the same or similar locations on the tree as introgression events we inferred with our DCT-BLT analysis (Figure S4). On the other hand, PhyloNet inferred an introgression event in clade 8 that is more ancient than that inferred by DCT-BLT (an introgression event between *Z. capensis* and the *Z. camerounensis*-*Z. nigranus* ancestor detected by DCT-BLT is pushed back to the *Z. camerounensis-Z. vittiger* ancestor by `PhyloNet`). Uncertainty over the precise history of introgression in clade 8 notwithstanding, PhyloNet is consistent with our DCT-BCT analysis and identifies introgression across the *Drosophila* phylogeny.

Finally, we asked whether the proportion of the genome that introgressed between putatively introgressing taxa (*γ*) varied with the timing of introgression events (Figure 4). Rather than timing introgression relative to when two hybridizing taxa shared a most recent common ancestor (which would require additional data, such as haplotype lengths of introgressed regions), we leveraged divergence time estimates across the drosophila phylogeny (Figure 1) and estimated when introgression events could have occured in time relative to the present (i.e., Mya). For this analysis, we focused on the 17 “best-supported” introgression events based on the criteria that more than 10% of the total triplets that could detect introgression between a given pair of taxa were significant according to both DCT and BLT (see solid red arrows in Figs. 3 and S3; Table 1). We estimated when these events occurred by taking the maximum, minimum, and midpoint times when the two branches that experienced introgression both coexisted in our dated phylogeny. We note that this approach results in imprecise time estimates, particularly for long branches in the phylogeny; however, it allowed us to test whether there was any obvious relationship between the proportion of the genome that introgressed (*γ*) and when those introgression events took place in the past. In one instance, the two branches that putatively experienced introgression did not overlap in time in our phylogeny. This situation could be explained by “ghost” introgression with unsampled or extinct lineages. For the 17 remaining introgression events, there was not a significant relationship between the midpoint estimate of timing of introgression (Mya) and *γ* (Spearman’s rank correlation: 0.43; *P* = 0.085; Figure 4). Our analyses therefore support introgression across the evolutionary history of *Drosophila*, with introgressing species pairs exchanging a similar fraction of the genome (range of average *γ* estimates = 0.013 - 0.237) regardless of whether those events were ancient or more recent.

**Figure 4.**
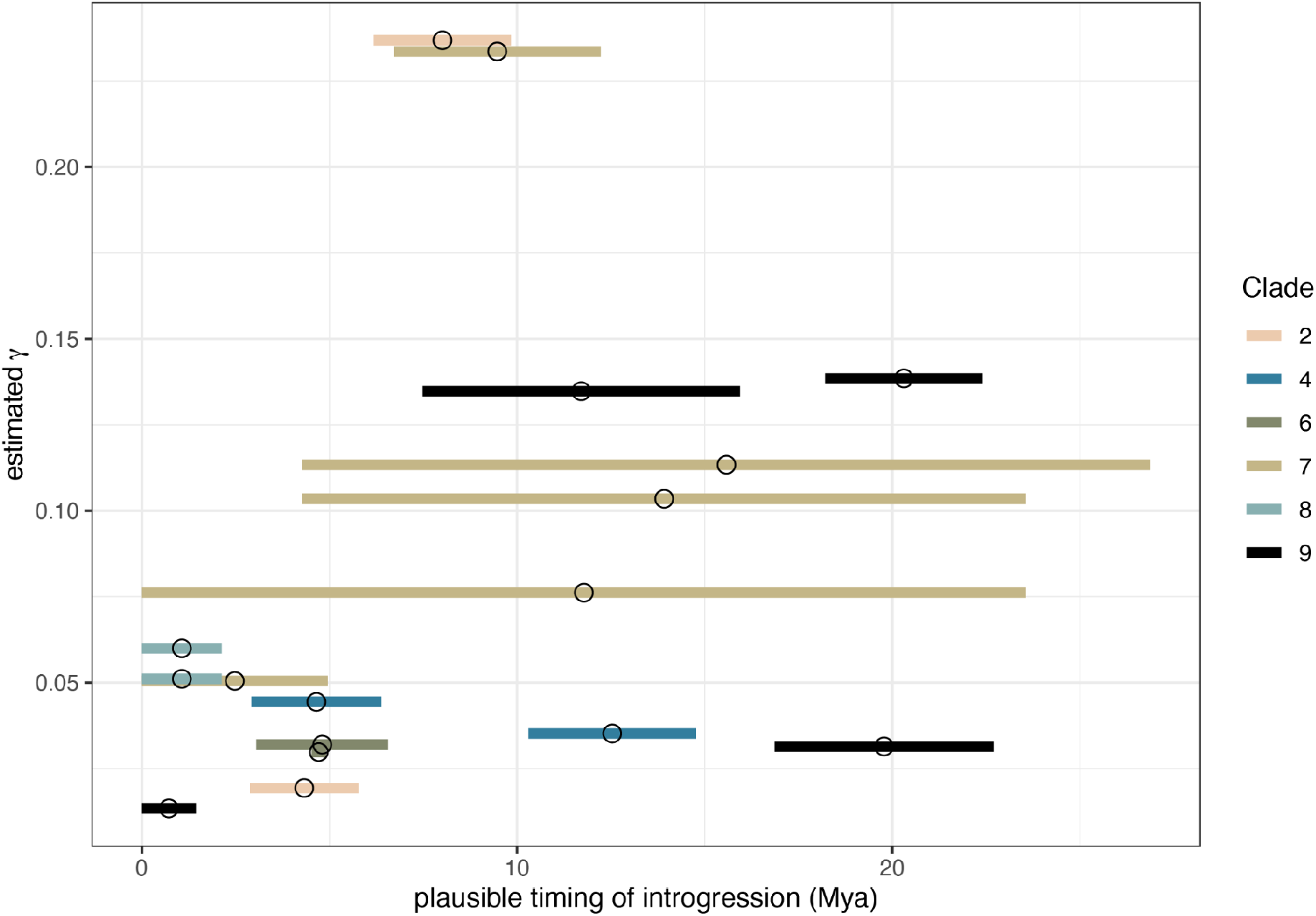
Time and fraction of the genome introgressing for the 17 best-supported introgression events across the *Drosophila* phylogeny. Each horizontal segment summarises one of the 17 introgression events highlighted in Figure 3 and is colored by clade. Segments span the times when the two putatively introgressing taxa both existed and are based on times inferred from the dating analysis summarised in Figure 1. Fraction of the genome that introgressed was estimated as the average f-branch statistic across all triplet comparisons that supported a given introgression event. Mya = million years ago.

## Discussion

### A time-calibrated tree of drosophilid evolution

*Drosophila*, as a genus, remains a premier model in genetics, ecology, and evolutionary biology. With over 1,600 species^42^, the genus has the potential to reveal why some groups are more speciose than others. Yet the phylogenetic relationships among the main groups in the genus have remained largely unresolved (reviewed in ^42^). Here we estimated a robust time-calibrated phylogeny for the whole genus using multilocus genomic data and calibrated it using a fossil record.

Our results confirm that the genus *Drosophila* is paraphyletic, with the genera *Zaprionus*, *Scaptomyza*, *Leucophenga*, and *Hirtodrosophila* each nested within the larger genus *Drosophila*. Consistent with the subdivisions previously proposed by refs. 60 and 44, clades 1-5 of our phylogeny contain species belonging to the subgenus *Sophophora*, and include species from the genus *Lordiphosa* (group A in Figure 1). Clades 6-9 of our phylogeny contain species belonging to the subgenus *Drosophila* (group B in Figure 1) and include species from the Hawaiian *Drosophila* and the subgenera *Siphlodora, Phloridosa* (synonymized with the subgenus *Drosophila^44^*, and genus *Zaprionus*. For more recent radiations within *Drosophila*, the topology we present is largely congruent with previous studies^42,51^ but two general observations are notable. First, our results confirm that *Lordiphosa* is closely related to the *saltans* and *willistoni* groups (clade 1) and part of the *Sophophora* subgenus (consistent with ref. ^61^). Second, we confirm that *Zaprionus* is related to the *cardini/qunaria/immigrans* group (consistent with refs. 42 and 60, but discordant with 43). Despite our well resolved phylogeny, comparisons with other studies emphasize the need to expand species sampling, especially given the potential to generate highly contiguous genomes at relatively low cost^62^.

Our results from divergence time analysis via `MCMCTree` suggest that the origin of *Drosophila* (including the subgenera *Sophophora* (group A) and *Drosophila* (group B)) occurred during the Eocene Epoch of the Paleogene, which is younger than estimates by ^60^ and ^43^, but older than estimates by ^49^. Different estimates of divergence times may be the result of different calibration information used, such as mutation rates, the time of formation of the Hawaiian Islands, and the fossil record. However, our comparison of various calibration schemes suggests that the choice of calibration information has a minor effect on `MCMCTree`’s age estimation (Figure S2). Additionally, credible intervals around our estimates tend to be notably narrower than in all of the aforementioned studies. In contrast to the previous studies, we used genome-scale multilocus data which would be expected to improve both the accuracy and precision of age estimates^63,64^.

On the other hand, we note that our analyses in `BEAST2` using the FBD model yielded significantly older ages (Figure S2) especially for deeper nodes and with markedly wider credible intervals suggesting origination of *Drosophila* lineage in the Late Cretaceous. These calibration inconsistencies may arise as a result of the poor fossil record within *Drosophila* (only *Scaptomyza dominicana* from Dominican amber) and selection of the oldest fossils for deeper radiations, which together can lead to overestimation of nodal ages under the FBD model^65^. Moreover, the poor convergence behavior we observed would also be expected to produce larger credible intervals.

### The extent of introgression in *Drosophila*

Access to genome-scale data has reinvigorated the study of hybridization and introgression^14^. We used genome-scale sequence data to provide the first systematic survey of introgression across the phylogeny of drosophilid flies. Our complementary—and conservative—approaches identified overlapping evidence for introgression within seven of the nine clades we analyzed (Figs. 3 and S3, Data S2). We conclude that at least 30 pairs of lineages have experienced introgression across *Drosophila*’s history (Table 1), though we note that other methods recover more introgression events (Data S2) and thus we cannot rule out the possibility that the true number is substantially higher. Moreover, we find that in many cases a substantial fraction of the genome is introgressed: our estimates indicate that numerous introgression events have altered gene tree topologies for >10% of the genome (Figs. 3 and S3, Table 1). Studies in contemporary *Drosophila* species suggest that selection may constrain the evolution of mixed ancestry, at least in naturally occurring^9,36,66^ and experimental admixed populations^67,68^. The results we have presented here used phylogenetic signals to show that introgression has nonetheless occurred and left a detectable signal within the genomes of many extant *Drosophila*.

In addition to providing an estimate of the extent of introgression, our results are informative about the timing of introgression among *Drosophila* lineages: the approaches we used to estimate the number of introgression events, and map them onto the phylogeny could potentially overestimate the timing of introgression if multiple independent more recent events are mistaken for one ancestral event. However, as described in the Results, both our PhyloNet analyses and a careful examination of our DCT-BLT results are most consistent with ancient introgression events in many cases. We also find evidence for very recent events, and although our analyses did not search for gene flow between sister taxa, previous studies of closely related species in *Drosophila* have revealed evidence of introgression^9,10,29,31,32^. Studies that have taken phylogenomic approaches to detect introgression in other taxa have also reported evidence for introgression between both “ancient” lineages (i.e., those that predate speciation events generating extant species) and extant species^8,12,18,19,21^. We conclude that introgression between *Drosophila* flies has similarly occurred throughout their evolutionary history.

Although the signal of introgression across our phylogeny provides evidence for widespread introgression in *Drosophila*, the evolutionary role of introgressed alleles remains to be tested. For example, the impact of hybridization and introgression on evolution can be diverse, from redistributing adaptive genetic variation^23,69,70^ to generating negative epistasis between alleles that have evolved in different genomic backgrounds (refs. 71–73; reviewed in refs. 15,16,74,75). The number of introgressed alleles that remain in a hybrid lineage depends on their selection coefficients^76–78^, their location in the genome (i.e., sex chromosomes vs. autosomes^79–81^), levels of divergence between the hybridizing species^9,82,83^, and recombination rates among loci^6,84^. Previous studies have, for example, shown that *Drosophila* hybrids often show maladaptive phenotypes^36,85–89^. Similarly, experimental hybrid swarms generated from two independent species pairs of *Drosophila* have shown that these populations can evolve to represent only one of their two parental species within as few as 10 generations, with the genome of one of their two parental species being rapidly purged from the populations^67^. These results show how hybrid *Drosophila* can be less fit than their parents, and further work is needed to determine the evolutionary effects, and the ecological context, of the introgression that we report here. However, our results suggest that not all introgressed material is deleterious in *Drosophila*, as we find that for some lineages a large fraction of the genome is introgressed (i.e. our *γ* estimates shown in Figs. 3 and S3 and Table 1). These results add to the growing body of literature that document a detectable phylogenetic signal of introgession left within the genomes of a wide range of species radiations that include *Drosophila*, other dipterans^90^, lepidopterans^8,84,91^, humans^5,92,93^, fungi^1,2^, and angiosperm plants^11,12^.

### Caveats and future directions

We estimated the number of events required to explain the introgression patterns across the tree and in some cases those events were recovered as relatively ancient. However, our approaches for mapping gene flow events onto the phylogeny was somewhat parsimonious in that it favors older events over repeated and recent introgressions (see Method Details), and thus may bias the age of introgression towards ancient events and underestimate the true number of pairs of lineages that have exchanged genetic material. For example, introgression events we inferred at deeper nodes in our phylogeny are often supported by only a subset of comparisons between species pairs that spanned those nodes (e.g. see “ancient” introgression events in clades 2, 7 and 9; Figure 3). It is also possible that some patterns we observe reflect scenarios where introgressed segments have persisted along some lineages but been purged along others. This phenomenon could also cause older gene flow between sister lineages, which should generally be undetectable according to the BLT and DCT methods, to instead appear as introgression between non-sister lineages that our methods can detect. Future work could seek to more precisely reveal the number and timing of gene flow events across this phylogeny, including more recent introgression events and gene flow between extant and extinct/unsampled lineages, a pattern referred to as “ghost” introgression^94,95^.

Our analyses also do not identify the precise alleles that have crossed species boundaries or reveal the manner in which these alleles may have affected fitness in the recipient population^74,75^. Genome alignments, complete annotations, and/or population level sampling across the genus are required to determine whether certain genes or functional categories of genes are more likely to cross species boundaries than others. More complete taxonomic sampling, combined with methodological advances for inferring the number and timing of introgression events in large phylogenies, will increase our ability to identify the specific timing and consequences of introgression across *Drosophila*.

### Conclusions

Speciation research has moved away from the debate of whether speciation can occur with gene flow to more quantitative tests of how much introgression occurs in nature, and how this introgression affects the fitness of individuals in the recipient population. Our well-resolved phylogeny and survey of introgression revealed that gene flow has been a relatively common feature across the evolutionary history of *Drosophila*. Yet, identifying the specific consequences of introgression on fitness and the evolution of species and entire radiations within *Drosophila* and other systems remains a major challenge. Future research could combine the power of phylogenomic inference with population-level sampling to detect segregating introgression between sister species to further our understanding of the amount, timing, and fitness consequences of admixture for diversification.

## METHOD DETAILS

### Genome assemblies and public data

Genome sequences used by this work were obtained from concurrent projects and public databases. Genome sequencing and assembly for 84 genomes is described in^62^. These data are available for download at NCBI BioProject PRJNA675888. For the remaining genomes: sequencing and assembly of 8 Hawaiian *Drosophila* were provided by E. Armstrong and D. Price, described in Armstrong et al. (in prep) and available at NCBI BioProject PRJNA593822; sequences and/or assemblies of five *nannoptera* group species were provided by M. Lang and V. Courtier-Orgogozo and are available at NCBI BioProject PRJNA611543; 44 were downloaded as assembled sequences from NCBI GenBank; *Z. sepsoides* and *D. neohypocausta* were sequenced as paired-end 150bp reads on Illumina HiSeq 4000 at UNC and assembled using SPAdes v3.11.1 with default parameters^103^; and 15 were generated by assembling short read sequences downloaded from NCBI SRA. For sets of unassembled short reads, we used ABySS v2.2.3^104^ with parameters “k=64” with paired-end reads (typically 100-150bp) to assemble the reads. Finally, outgroup genome sequences (*A. gambiae*, *M. domestica*, *L. trifolii*, *C. hians*, and *E. gracilis*) were obtained from NCBI GenBank. See Data S3 for a full list of samples, strain information, accessions, and associated publications.

### Orthology Inference

We identified single-copy orthologous genes in each genome using BUSCO (Benchmarking Universal Single-Copy Orthologs; v3.1.0^98^). BUSCO was run with orthologs from the Diptera set in OrthoDB v.9 (odb9) using default parameters. For each species, all BUSCOs found in a single copy were used for phylogenetic analysis.

### Assignment of BUSCO genes to Muller elements for *obscura* group species

Each of the BUSCO genes identified as single-copy in each of the group 12 (*obscura* group: *D. affinis*, *D. athabasca*, *D. azteca*, *D. bifasciata*, *D. guanche*, *D. lowei*, *D. miranda*, *D. obscura*, *D. persimilis*, *D. pseudoobscura*, *D. subobscura*) genome assemblies was assigned to one of the six Muller elements (elements A-F). For *D. athabasca*, *D. bifasciata*, *D. lowei*, *D. miranda*, *D. pseudoobscura*, and *D. subobscura*, contig/scaffold associations with chromosomes and/or Muller elements were simply obtained from NCBI GenBank assembly report tables. For the remaining genomes (*D. affinis*, *D. azteca*, *D. guanche*, *D. obscura*, *D. persimilis*), we used whole-genome alignments to infer the Muller element associated with each contig or scaffold. Using the Progressive Cactus^105^ software, each remaining genome was aligned to a closely related reference genome (*D. affinis* - *D. athabasca; D. azteca* - *D. athabasca; D. guanche* - *D. subobscura; D. obscura* - *D. bifasciata; D. persimilis* - *D. miranda*) with a similar karyotype^54,106^. Using the reference genomes as backbones, each remaining genome was then scaffolded, with Ragout^107^. The scaffolds allowed us to annotate each contig in the remaining genomes with Muller element information from the reference genomes (see Data S4). BUSCO genes on unplaced contigs were ignored.

### Phylogenetic reconstruction

Every DNA BUSCO locus was aligned with MAFFT v7.427^100^ using the L-INS-i method. We removed sites that had fewer than three non-gap characters from the resulting multiple sequence alignments (MSAs). These trimmed MSAs were concatenated to form a supermatrix. To assess the quality of the assembled supermatrices we computed pairwise completeness scores in AliStat^108^ (Figure S5) . We inferred a maximum likelihood (ML) phylogenetic tree from the supermatrix (a.k.a. concatenated alignment) using IQ-TREE v1.6.5^99^, and treated the supermatrix as a single partition. IQ-TREE was run under GTR+I+G substitution model, as inference under any other substitution model will not necessarily lead to better accuracy of tree topology estimation^109^. To estimate the support for each node in this tree, we used three different reliability measures. We did 1,000 ultrafast bootstrap (UFBoot) replicates^110^ and additionally performed an approximate likelihood ratio test with the nonparametric Shimodaira–Hasegawa correction (SH-aLRT) and a Bayesian-like transformation of aLRT^111^. We used the ML gene trees obtained by IQ-TREE with a GTR+I+G substitution model for tree inference in ASTRAL^96^. For the estimated ASTRAL tree we calculated the support of each node using local posterior probabilities (LPP)^96^. Also, we created a gene tree set by removing taxa with outlier branch lengths that were potentially produced by misaligned regions and/or incorrect orthology inference in TreeShrink^45^ under default parameters. This analysis resulted in a small fraction of branches removed from our gene tree set (<5.5%)

We did two additional analyses to verify the robustness of our topology inference. First, we inferred an ML tree using WAG+I+G substitution model from the protein supermatrix obtained from concatenation of protein BUSCO MSAs. MSAs based on amino acid sequences have been shown to have superior accuracy to DNA MSAs for distantly related species^112^. Second, to verify that long branch attraction did not distort our tree topology, we inferred an ML tree under a GTR+I+G substitution model using a different set of outgroup species from the DNA supermatrix. Specifically, instead of distantly related *Anopheles gambiae*, we used *Musca domestica*, *Liriomyza trifolii*, *Curricula hians* and *Ephydra gracilis* together as our outgroup species.

### Phylogenetic Support Analysis via Quartet Sampling

We used quartet sampling (QS) as an additional approach to estimate phylogenetic support^11^. Briefly, QS provides three scores for internal nodes: (*i*) quartet concordance (QC), which gives an estimate of how sampled quartet topologies agree with the putative species tree; (*ii*) quartet differential (QD) which estimates frequency skewness of the discordant quartet topologies, and can be indicative of introgression if a skewed frequency observed, and (*iii*) quartet informativeness (QI) which quantifies how informative sampled quartets are by comparing likelihood scores of alternative quartet topologies. Finally, QS provides a score for terminal nodes, quartet fidelity (QF), which measures a taxon “rogueness”. We did QS analysis using the DNA BUSCO supermatrix described above, specifying an IQ-TREE engine for quartet likelihood calculations with 100 replicates (i.e. number of quartet draws per focal branch).

### Fossil Dating

#### MCMCTREE

We implemented the Bayesian algorithm of MCMCTree v4.9h^46^ with approximate likelihood computation to estimate divergence times within *Drosophila* using several calibration schemes (Data S1). First, we estimated branch lengths by ML and then the gradient and Hessian matrices around these ML estimates in MCMCTree using the DNA supermatrix and species tree topology estimated by IQ-TREE. Because large amounts of sequence data are not essential for accurate fossil calibration^113^, we performed dating analysis using a random sample of 1,000 MSA loci (out of 2,791) for the sake of computational efficiency. Thus, for this analysis the supermatrix was generated by concatenating 1,000 randomly selected gene-specific MSAs. Using fewer loci (10 and 100) for fossil calibration did not drastically affect nodal age estimation (Figure S1). We removed sites that had less than 80 non-gap characters from all these supermatrices. Second, we used the gradient and Hessian matrix, which constructs an approximate likelihood function by Taylor expansion^114^, to perform fossil calibration in MCMC framework. For this step we specified a GTR+G substitution model with four gamma categories; birth, death and sampling parameters of 1, 1 and 0.1, respectively. To model rate variation we used an uncorrelated relaxed clock. To ensure convergence, the analysis was run ten times independently for 8 × 10^6^ generations (first 10^6^ generations were discarded as burn-in), logging every 1,000 generations. We used the R package MCMCtreeR^101^ to visualize the calibrated tree.

#### BEAST 2

Additionally we performed fossil calibration using the Fossilized Birth-Death (FBD) process^47^ as implemented in the Bayesian framework of BEAST 2.6.3^48^. For scalability purposes, we randomly selected 1,000 loci and then partitioned them into 10 supermatrices each consistent of 100 different MSAs. Each of these 10 datasets was treated as a single partition in the downstream analyses. Additionally, we removed sites that had less than 128 non-gap characters from all these supermatrices. To perform fossil calibration, we used a GTR+G model with four gmamma categories, and an optimized relaxed clock^115^ was used to model rate variation. For the FBD prior we specified an initial origin value of 230 Mya (which corresponds to the age of oldest known dipteran fossil *Grauvogelia*), and the tree likelihood was conditioned on the proportion of species sampled at present (*ρ* = 0.1). The remaining priors were set to their defaults. In order to directly compare divergence time estimation between BEAST 2 and MCMCTree, we used the same fixed IQ-TREE species tree topology with several exceptions. First, we did not fix the phylogenetic positions of contemporary *Scaptomyza* species and fossil taxon *Scaptomyza dominicana* within its monophyletic group. Second, we did not constrain relationships of outgroup species *L. varia, C. costata, S. lebanonensis* including fossil taxon *Electrophortica succini*. Two additional fossils, *Oligophryne* and *Phytomyzites*, were specified for Drosophilidae stem. Furthermore, to accomodate uncertainty of fossil dates we incorporated age ranges for several fossils (Data S1). For each of the 10 datasets we ran 2 independent MCMC chains for 6 × 10^8^ generations with sampling frequency of 10,000 for each model parameter. Additionally, we performed sampling from the prior distribution only. Convergence was assessed using ESS in Tracer^102^. Divergence times were generated by taking means of posterior nodal ages discarding 25% of the sampled trees as burn-in in TreeAnnotator for each dataset. To drastically improve computational efficiency of likelihood calculations in all BEAST 2 analyses we used the program in conjunction with BEAGLE library^97^ that enables GPU utilization.

### Inferring Introgression Across the Tree

#### Gene tree-based methods

In order to detect patterns of introgression we used three different methods that rely on the topologies of gene trees, and the distributions of their corresponding branch lengths, for triplets of species. If the true species tree is ((A, B), C), these tests are able to detect cases of introgression between A and C, or between B and C. These include two of the methods that we devised for this study, and which use complementary pieces of information—the counts of loci supporting either discordant topology, and the branch-length distributions of gene trees supporting these topologies, respectively—to test an introgression-free null model.

The first method we developed was the discordant-count test (DCT), which compares the number of genes supporting each of the two possible discordant gene trees: ((A, C), B) or (A, (B, C)), similar in principle to the delta statistic from^116^. Genes may support the two discordant topologies (denoted T_1_ and T_2_) in the presence of ILS and/or in the presence of introgression. In the absence of ancestral population structure, gene genealogies from loci experiencing ILS will show either topology with equal probability; ILS alone is not expected to bias the count towards one of the topologies. In the presence of introgression, one of the two topologies will be more frequent than the other because the pair of species experiencing gene flow will be sister lineages at all introgressed loci (illustrated in Figure 2). For example, if there is introgression between A and C, there will be an excess of gene trees with the ((A, C), B) topology. The DCT identifies pairs of species that may have experienced introgression by performing a χ^2^ goodness-of-fit test on the gene tree count values for a species triplet to determine whether their proportions significantly deviate from 0.5, the expected proportion for each gene genealogy under ILS. We used this test on all triplets extracted from BUSCO gene trees within each clade, and the resulting *P*-values were then corrected for multiple testing using the Benjamini-Hochberg procedure with a false discovery rate (FDR) cutoff of 0.05. We note that these tests are not independent since different triplets may contain overlapping taxa. Thus, while our correction results in more conservative tests^57^, the inferred FDRs may be somewhat inaccurate.

Second, we devised a branch-length test (BLT) to identify cases of introgression (illustrated in Figure 2). This test examines branch lengths to estimate the age of the most recent coalescence event (measured in substitutions per site). Introgression should result in more recent coalescences than expected under the concordant topology with complete lineage sorting, while ILS shows older coalescence events^90^. Importantly, ILS alone is not expected to result in different coalescence times between the two discordant topologies, and this forms the null hypothesis for the BLT. For a given triplet, for each gene tree we calculated the distance *d* (a proxy for the divergence time between sister taxa) by averaging the external branch lengths leading to the two sister taxa under that gene tree topology. We calculated *d* for each gene tree and denote values of *d* from the first discordant topology *d*_T1_ and those from the second discordant topology *d*_T2_. We then compared the distributions of *d*_T1_ and *d*_T2_ using a Mann-Whitney *U* test. Under ILS alone the expectation is that *d*_T1_ = *d*_T2_, while in the presence of introgression *d*_T1_ < *d*_T2_ (suggesting introgression consistent with discordant topology T1) or *d*_T1_ > *d*_T2_ (suggesting introgression with consistent with topology discordant T_2_). The BLT is conceptually similar to the D3 test^117^, which transforms the values of *d*_T1_ and *d*_T2_ in a manner similar to the *D* statistic for detecting introgression^92^. As with the DCT, we performed the BLT on all triplets within a clade and used a Benjamini-Hochberg correction with a false discovery rate cutoff (FDR) of 0.05. We note that both the DCT and BLT will be conservative in cases where, for a triplet ((A,B), C), there is introgression between A and C as well as B and C, with the extreme case of equal rates of introgression for both species pairs resulting in a complete loss of power.

Finally, we used QuIBL^8^, an analysis of branch-length distribution across gene trees to infer putative introgression patterns. Briefly, under coalescent theory internal branches of rooted gene trees for a set of 3 taxa (triplet) can be viewed as a mixture of two distributions: one that generates branch lengths under ILS, and the other under introgression/speciation. Thus, the estimated mixing proportions (π_1_ for ILS and π_2_ for introgression/speciation; π_1_ +π_2_ = 1) of those distribution components show which fraction of the gene trees were generated through ILS or non-ILS processes. For a given triplet, QuIBL computes the proportion of gene trees that support the three alternative topologies. Then for every alternative topology QuIBL estimates mixing proportions along with other relevant parameters via Expectation-Maximization and computes Bayesian Information Criterion (BIC) scores for ILS-only and introgression models. For concordant topologies elevated values of π_2_ are expected whereas for discordant ones π_2_ values significantly greater than zero are indicative of introgression. To identify significant cases of introgression here we used a cutoff of ΔBIC < −30 as in^8^. We ran QuIBL on every triplet individually under default parameters with the number of steps (the _numsteps_ parameter) set to 50 and using *Anopheles gambiae* for triplet rooting; the branch length between *A. gambiae* and the triplet is not used for any of QuIBL’s calculations.

We note that the DCT and BLT methods are potentially impacted by ancestral population structure: if the lineages leading to B and C were in subpopulations that were more likely to interbreed in the ancestral population, then the ((B, C), A) topology might be expected to be more prevalent than ((A, C), B), along with a shorter time back to the first coalescence. However, it is unclear how much of a concern ancestral population structure should be for this analysis, as it seems less likely that it would be a pair of lineages that diverged first (i.e. A and C or B and C) that interbred more frequently in the ancestral population instead of the two lineages that went on to be sister taxa (i.e. A and B). Nonetheless, plausible scenarios of ancestral structure supporting one discordant topology over the other can be devised (e.g. ref ^118^). We therefore conducted a more stringent version of our DCT-BLT combined test that requires the average distance between the two introgressing taxa (when examining gene trees with the discordant topology consistent with introgression) to be less than that between the two sister species (when examining gene trees with the concordant topology). Such a pattern is consistent with introgression between non-sister species, which must occur more recently than the species split and therefore causing more recent coalescence events, but not with ancestral structure which will still result in older coalescence times for discordant trees than the concordant trees (because structure in the ancestral population is only a factor in the case of ILS). Note that this test is expected to be especially conservative because ILS, which for many triplets accounts for a sizable fraction of our discordant gene trees, will push the coalescent times for all discordant topologies back further in time.

We also examined the effect of evolutionary rate heterogeneity measured in branch-specific *d*_N_/*d*_S_ values on introgression detection. To that end, we generated codon alignments for each BUSCO locus using TranslatorX^119^ and then calculated *d*_N_/*d*_S_ ratios for each gene tree in PAML^46^ within each clade using a free-ratios branch model that assumes independent *d*_N_/*d*_S_ for each gene tree branch. Then, we evaluated the distribution of *d*_N_/*d*_S_ ratios across all gene trees to determine the 95^th^ percentile value of *d*_N_/*d*_S_. Thus, we repeated our DCT/BLT analyses for each triplet after excluding every gene tree that had at least one branch with *d*_N_/*d*_S_ > 0.53. Note, branches with *d*_N_/*d*_S_ values where *d*_S_< 0.001 or >5 were deemed unreliable and thus were excluded from calculation of a critical value or from downstream filtering. Additionally, we performed random filtering of gene trees to see if this procedure would have a similar impact on downstream introgression-detection as did our *d*_N_/*d*_S_ filter. First, we estimated the distribution of proportions of gene trees retained for each triplet after applying the *d*_N_/*d*_S_ filter. Then, for a given triplet, we randomly drew a number of genes to remove from the aforementioned distribution, and then applied our DCT-BLT method to this triplet after removing the selected number of genes. This process was repeated for each triplet tested in our main analysis to generate a randomly filtered set of DCT-BLT results for each of our 9 clades. We then repeated this entire process 1000 times and noted the average fraction of DCT-BLT results remaining significant after randomly filtering genes.

Our DCT-BLT test assumes that there is no recombination within loci and complete inter-locus independence—these assumptions are commonly made by introgression inference methods^10,120,121^. We note that intra-locus recombination may interfere with the signatures of introgression by reducing discordant topology counts (because even loci experiencing introgression will have non-introgressed segments), and similarly diluting branch-length signatures of introgression, thereby reducing the sensitivity of our DCT-BLT approach. Nevertheless, site-pattern-based approaches (e.g. HyDe, see below) are not affected by intra-locus recombination as they evaluate each site in an MSA independently.

#### Site-pattern-based detection of introgression

Signatures of introgression can be identified by investigating fractions of certain site patterns within MSAs of species quartets. One of the most widely used methods is based on the counts of ABBA-BABA site patterns (aka., Patterson’s D statistic^122^). Here we used the hybridization model implemented in HyDe^52^ that implements an alternative invariant-based statistic to test introgression and estimate the fraction of the introgressed genome (*γ*). We ran HyDe analysis on each of the 9 clades using the entire supermatrix and in each case selected the quartet’s outgroup from a sister clade. Additionally, to examine effects of outgroup choice, we ran HyDe analyses with a more distantly related outgroup, *Anopheles gambiae* for all clades. The resulting *P*-values for each quartet were corrected for multiple testing using the Bonferroni method. To investigate an individual contribution of each BUSCO locus to introgression, we additionally ran HyDe using BUSCO MSAs with *Anopheles gambiae* outgroup. We note, however, in this case HyDe’s power to detect introgression will be reduced, especially for short MSAs with <10,000 sites^52^. A complete summary for each BUSCO locus including introgression results from locus-specific HyDe and BLT/DCT analyses is included in Data S5.

#### Placing introgression events on the phylogeny

All the aforementioned methods can infer multiple correlated signatures of introgression especially when triplets/quartets share the same taxa. Thus it can be difficult to interpret these interdependent results. To alleviate this problem,^20^ devised a simple heuristic metric called *f*-branch to disentangle and map introgression events detected in multiple correlated species pairs onto the tree. In the original formulation, *f*-branch examines multiple *f_4_* statistics measured for each species pair and that quantify *γ*, the proportion of introgressed material for that pair. However, the calculation of the *f*_4_ statistic requires allele frequency measures within each sampled species. Thus, to calculate *f*-branch statistic, instead of *f*_4_ we used the introgression proportion derived from DCT/BLT as follows: 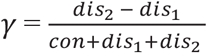, where *con*, *dis_1_* and *dis_2_* represent concordant and discordant counts of gene trees and *dis_1_* < *dis_2_*. To compute *f*-branch statistic from DCT/BLT’s *γ* estimates and to visualize the results within each clade we used the `Dsuite` python package^57^.

`Dsuite` outputs a matrix of *γ* estimates that have been partially collapsed: on one axis of this matrix signals of introgression can appear on ancestral branches, but on the other axis only extant branches are shown. Thus, we manually further collapsed these signatures by parsimoniously assuming that if some lineage A showed evidence of introgression with multiple descendants of some other lineage B that is not ancestral to A, then we considered this to be caused by a single introgression event between A and B. Note that we did not require all descendants of lineage B to share this signature of introgression, and thus this approach could potentially undercount the number of introgression events and overestimate their ages.

#### Phylogenetic networks

Introgression generates instances of reticulate evolution such that purely bifurcating trees cannot adequately represent evolutionary history; phylogenetic networks have been shown to provide a better fit to describe these patterns^123,124^. We used PhyloNet^58,59^ to calculate likelihood scores for networks generated by placing a single reticulation event (node) in an exhaustive manner, i.e. connecting all possible branch pairs within a clade. Because full likelihood calculations with PhyloNet can be prohibitively slow for large networks, for each of clades 1 through 9 we selected a subsample of 10 species in a manner that preserves the overall species tree topology. No subsampling was performed for clade 3 which has fewer than 10 species. Using these subsampled clade topologies, we formed all possible network topologies having a single reticulation node (with the exception of networks having reticulation nodes connecting sister taxa). Because PhyloNet takes gene trees as input, for each clade we subsampled each gene tree to include only the subset of 10 species selected for the PhyloNet analysis (or all species in the case of clade 3); any gene trees missing at least one of these species were omitted from the analysis. Finally, we used the GalGTProb program^125^ of the PhyloNet suite to obtain a likelihood score for each network topology for each clade. We report networks with the highest likelihood scores.

## Data and code availability

The data and code produced during this study are publically available on GitHub (https://github.com/SchriderLab/drosophila_phylogeny) and FigShare (dx.doi.org/10.6084/m9.figshare.13264697). Whole genome sequencing data generated for this study are available on NCBI (BioProject PRJNA675888, BioProject PRJNA593822, and BioProject PRJNA611543).

## Acknowledgements

We thank M. Hahn, M. Turelli, A. Yassin, and M. Matschiner for helpful feedback on a previous draft, and M. Hibbins for sharing simulated gene trees. AS and DRS were supported by the NIH under award nos. R00HG008696 and R35GM138286. BYK was supported by the NIH under award no. F32GM135998. JW was supported by the NIH under award no. K01DK119582. DP, AAC were supported by NSF Dimensions of Biodiversity award 1737752. DRM and ERRD were supported by NIH award R01GM121750. The funders had no role in study design, data collection and analysis, decision to publish, or preparation of the manuscript.

## Competing interests

The authors declare that there is no conflict of interest

## Notes

### Competing Interest Statement

The authors have declared no competing interest.

