## Supplemental Information for "Widespread introgression across a phylogeny of 155 *Drosophila* genomes"

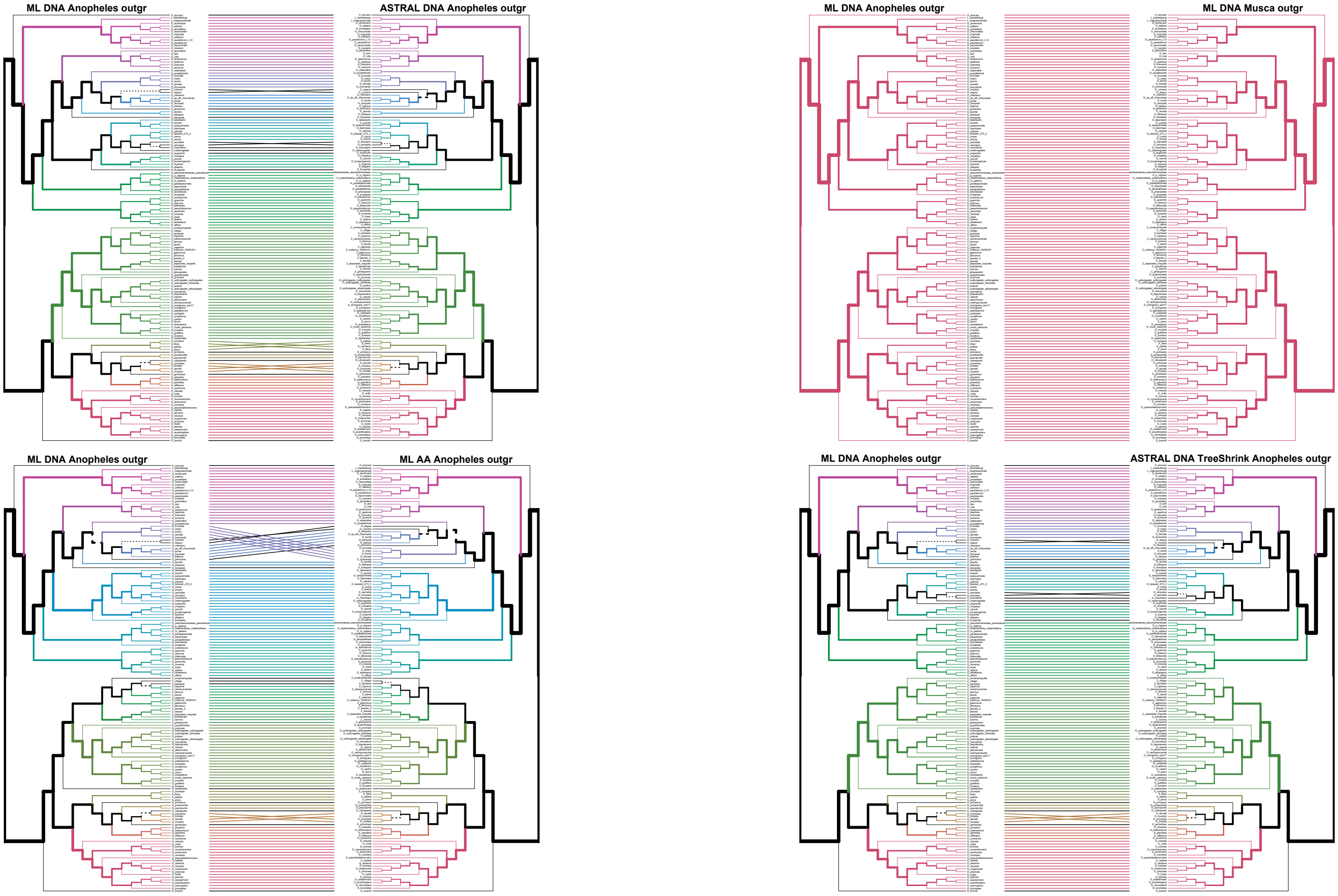


**Figure S1. Topological comparisons between trees inferred by different methods, related to Figure 1.** Black paths on the phylogenetic trees lead to dashed branches that indicate topological incongruencies.

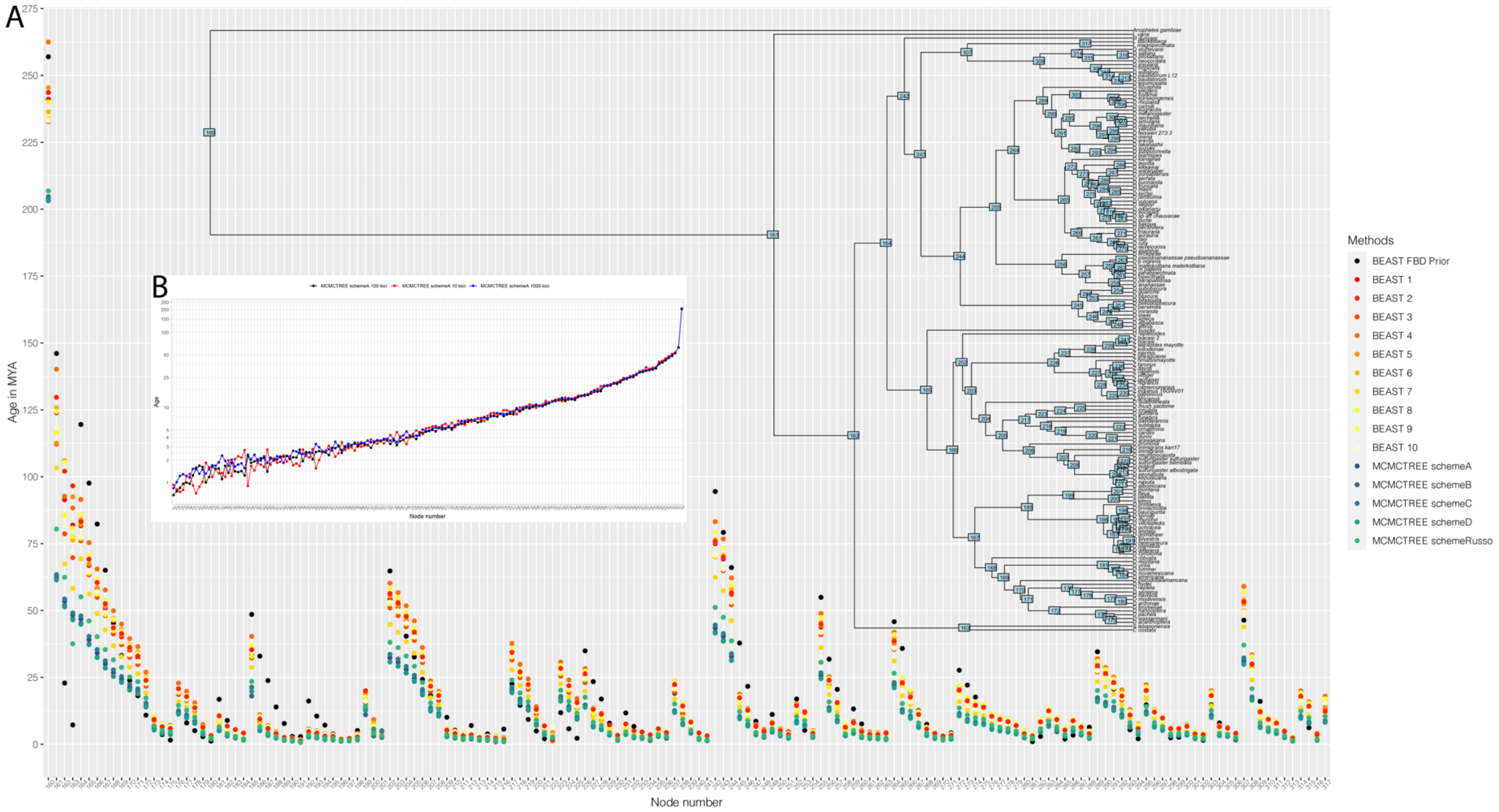


**Figure S2. Divergence time estimation, related to Figure 1.**
(A) Age estimates produced by different calibration schemes and methods. *x*-axis indicates node numbers. The *y*-axis represents the inferred age in MYA.
(B) Comparison between nodal calibrations based on different numbers of loci using scheme A. Nodes are arranged on the *x*-axis according to scheme A from the youngest to the oldest. The *y*-axis represents the log of inferred age in MYA.

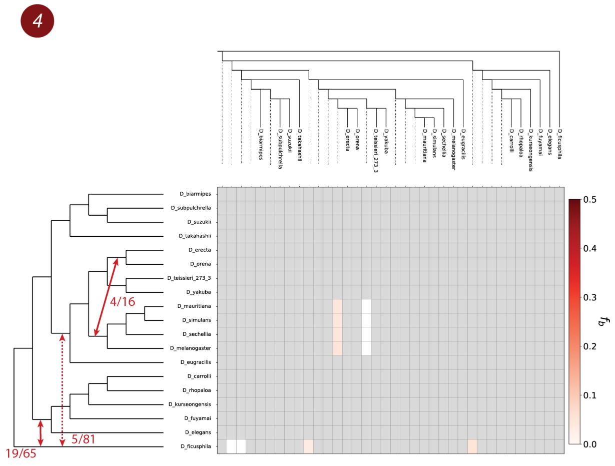
4
4/16
5/81
19/65
D_biarmipes
D_subpulchrella
D_suzukii
D_takahashii
D_erecta
D_orena
D_teissieri_273_3
D_yakuba
D_mauritiana
D_simulans
D_sechellia
D_melanogaster
D_eugracilis
D_carrolli
D_rhopaloa
D_kurseongensis
D_fuyamai
D_elegans
D_ficusphila
$f_b$
0.0
0.1
0.2
0.3
0.4
0.5

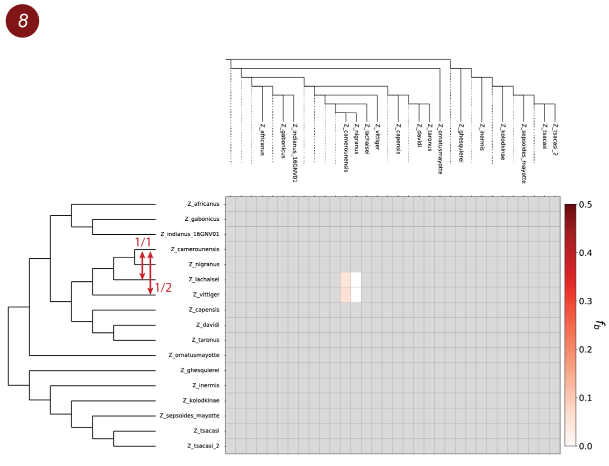
8
1/1
1/2
Z_africanus
Z_gabonicus
Z_indianus_16GNV01
Z_camerounensis
Z_nigranus
Z_lachaisei
Z_vittiger
Z_capensis
Z_davidi
Z_taronus
Z_ornatusmayotte
Z_ghesquierei
Z_inermis
Z_kolodkinae
Z_sepsoides_mayotte
Z_tsacasi
Z_tsacasi_2
$f_b$
0.0
0.1
0.2
0.3
0.4
0.5

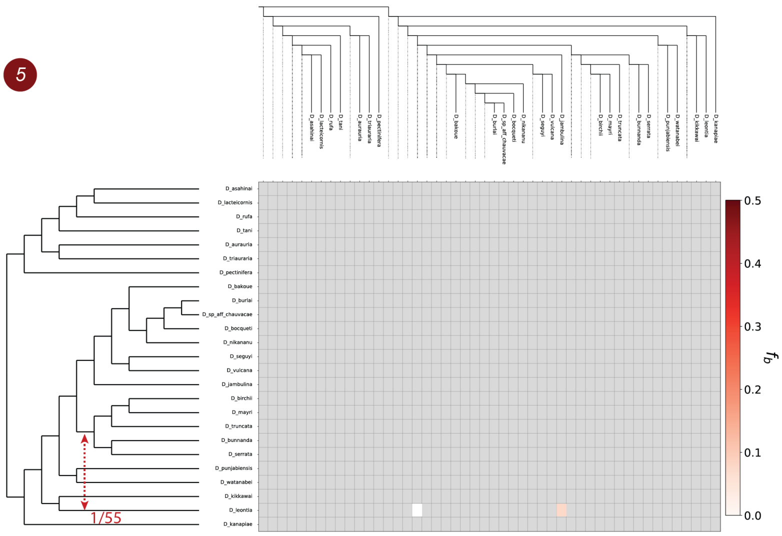
5
1/55
$f_b$
0.0
0.1
0.2
0.3
0.4
0.5

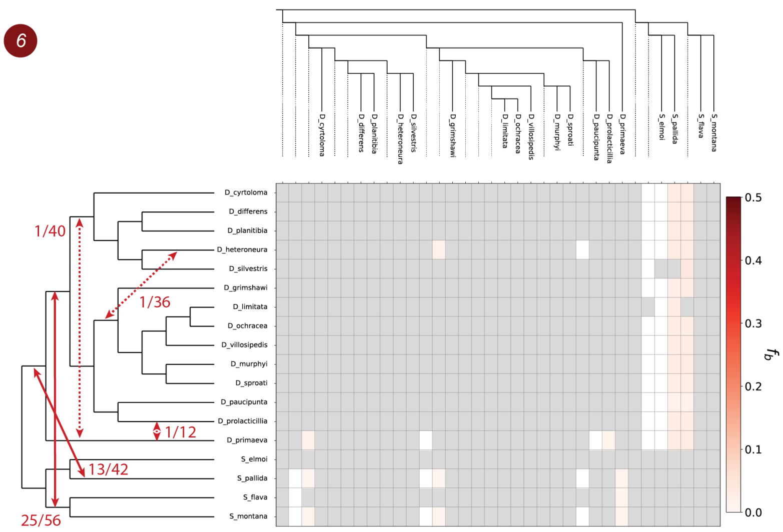
6
1/40
1/36
1/12
13/42
25/56
D_cyrtoloma
D_differens
D_planitibia
D_heteroneura
D_silvestris
D_grimshawi
D_limitata
D_ochracea
D_villosipedis
D_murphyi
D_sproati
D_paucipunta
D_prolacticillia
D_primaeva
S_elmoi
S_pallida
S_flava
S_montana
$f_b$
0.0
0.1
0.2
0.3
0.4
0.5

**Figure S3. Patterns of introgression inferred for the monophyletic clades 4–6 and 8, related to Figure 3.** The matrix shows inferred introgression proportions as estimated from gene tree counts for the introgressed species pairs (STAR Methods), and then mapped to internal branches using the *f*-branch method. The expanded tree at the top of each matrix shows both terminal and ancestral branches. The tree on the left side of each matrix represents species relationships with mapped introgression events (red arrows) derived from the corresponding *f*-branch matrix (STAR Methods). The fractions next to each arrow represent the number of triplets that support a specific introgression event by both DCT and BLT divided by the total number of triplets that could have detected the introgression event. Dashed arrows represent introgression events with low support (triplet support ratio < 10%). Clades 1 and 3 are not shown because no introgression was found in these clades according to the DCT-BLT combined test.

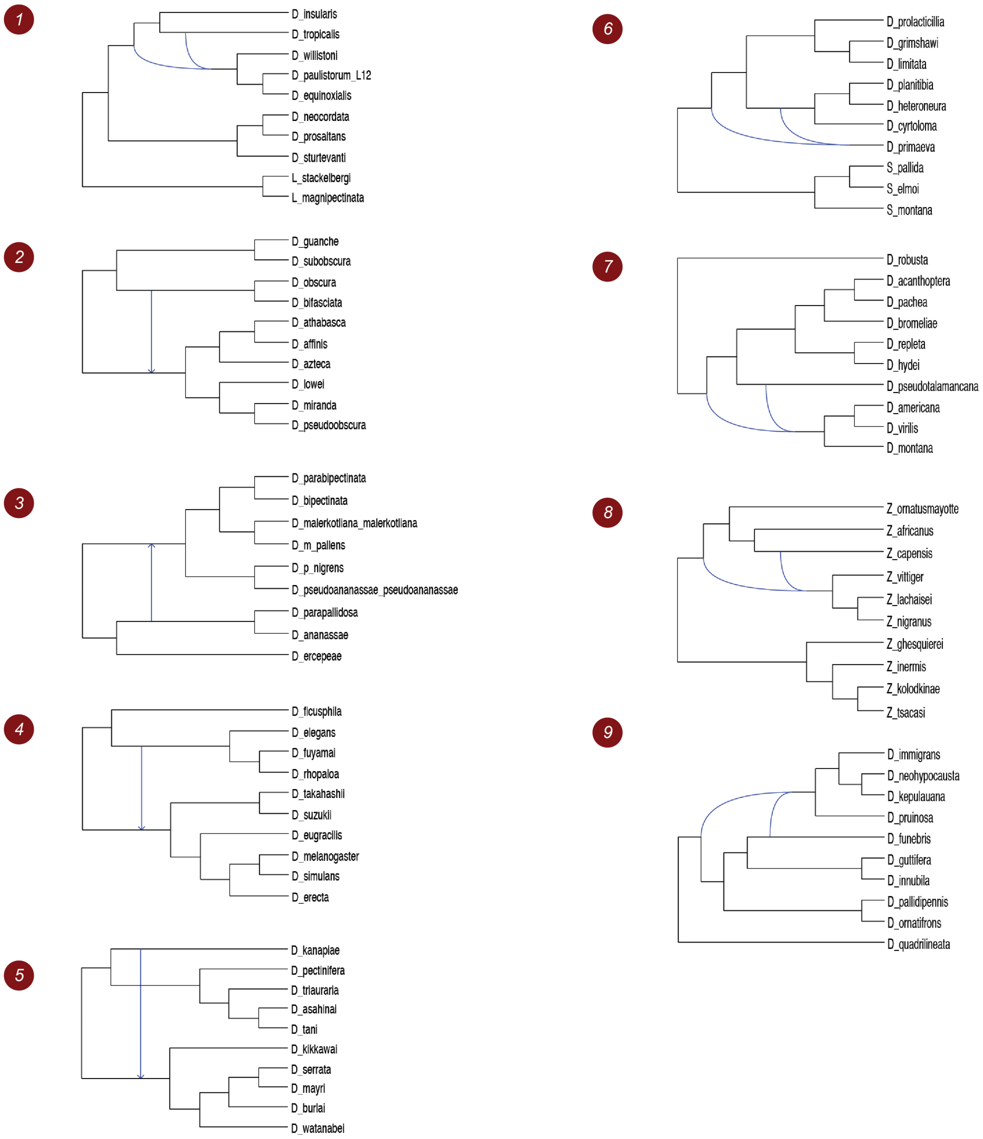


**Figure S4. Patterns of introgression inferred for the monophyletic clades 1–9 using PhyloNet, related to Figure 3.** For each clade we show the topology of the phylogenetic network with the highest likelihood. Blue branches denote reticulations.

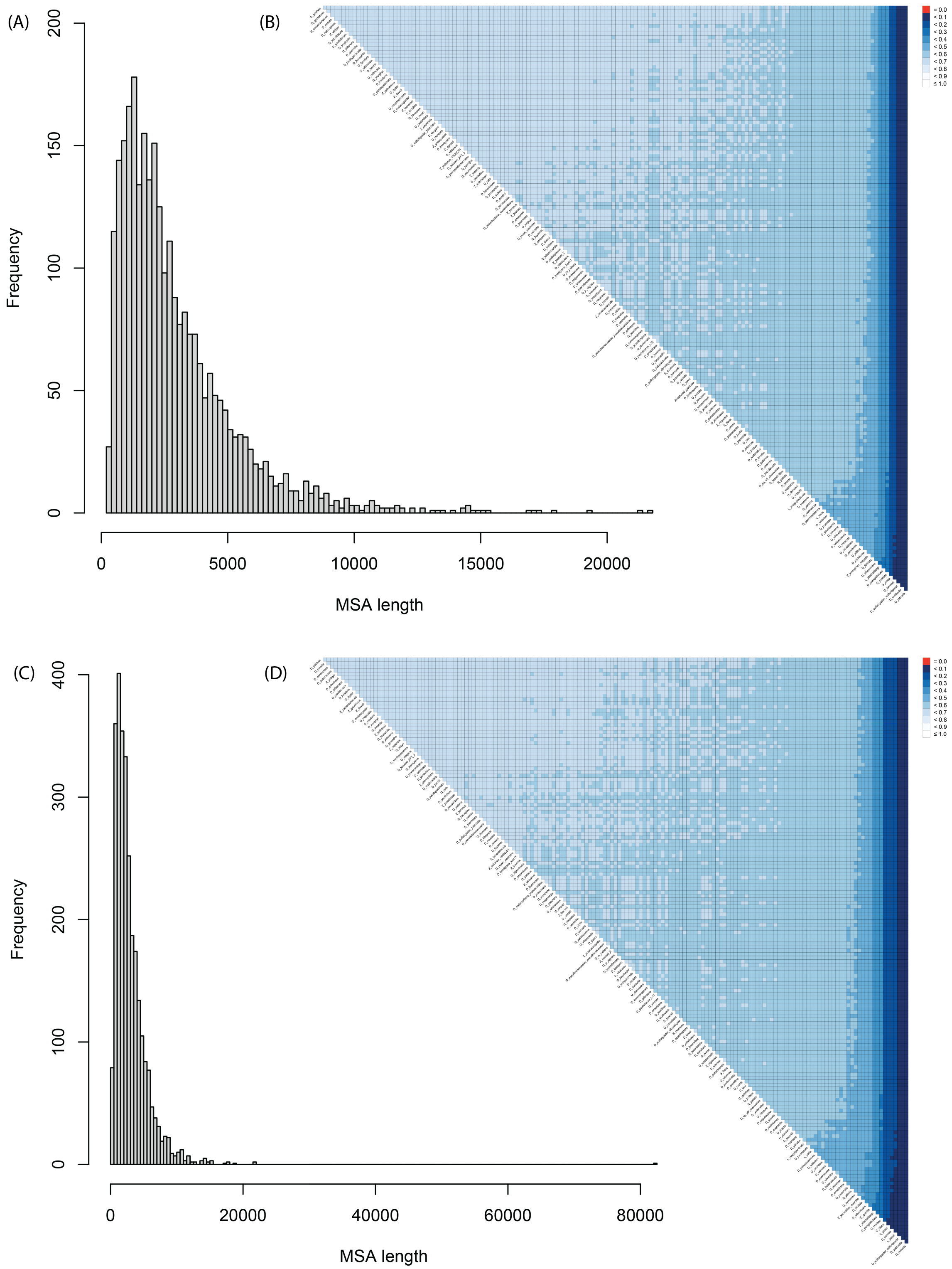


**Figure S5. Supermatrix completeness scores and distribution of BUSCO MSA lengths, related to Figure 1.** Matrices show pairwise completeness metric values computed by AliStat (the fraction of sites for which both sequences have a completely specified character). Larger values indicate higher levels of MSA completeness. MSA length distributions and AliStats matrices were generated for datasets where *Anopheles gambiae* (panels A and B) and *Musca domestica* (panels C and D) were used as outgroups.
